# Saccades and pre-saccadic stimulus repetition alter cortical network topology and dynamics: evidence from EEG and graph theoretical analysis

**DOI:** 10.1101/2021.06.08.447611

**Authors:** Amirhossein Ghaderi, Matthias Niemeier, John Douglas Crawford

## Abstract

Parietal and frontal cortex are involved in saccade generation, but their output signals also modify visual signals throughout cortex. These signals produce well-documented behavioral phenomena (saccades, saccadic suppression, various perisaccadic perceptual distortions) but their underlying influence on cortical network dynamics is not known. Here, we combined electroencephalography (EEG) with frequency-dependent source localization and graph theory analysis (GTA) to understand how saccades and pre-saccadic visual stimuli interactively alter cortical network dynamics in humans. 21 participants viewed series of 1-3 vertical or horizontal grids, followed by grid with the opposite orientation just before a horizontal saccade or continued fixation. EEG signals from the presaccadic interval (cue + 200ms, or equivalent fixation period) were used for source localization. Source localization (saccade – fixation) identified bilateral dorsomedial frontoparietal activity across frequency bands, whereas stimulus repetition produced band-specific modulations in left prefrontal, posterior parietal, and central-superior frontal and/or parietal cortex, with significant saccade-repetition interactions in frontal and parietal regions. GTA analysis revealed a saccade-specific functional network with major hubs in inferior parietal cortex (alpha) and the frontal eye fields (beta), and major saccade-repetition interactions in left prefrontal (theta) and supramarginal gyrus (gamma). Overall, quantitative measures of whole-brain network topology and dynamics (segregation, integration, synchronization, complexity) were enhanced during the presaccadic interval, but repetition interactions reduced synchronization and complexity. These results show that presaccadic signals have widespread, coherent influence on cortical network dynamics, likely responsible for both saccade production and the perceptual phenomena associated with saccades.

**Highlights:**

- Source localization & graph theory were used to analyze presaccadic EEG signals
- Presaccadic signals produced band-specific modulations/hubs in parietofrontal cortex
- Frontal/parietal eye fields showed extensive functional connectivity across all lobes
- Presaccadic stimulus repetition further modulated functional network connectivity
- Saccades and repetition both influenced network clustering, integration, & complexity

## Introduction

Every day we perform thousands of saccades for purposes ranging from reading to looking at a landscape (Kleiser, Seitz, and Krekelberg 2004). During each saccade, the eye rapidly moves to a new fixation point in order to maximize foveal vision (Bahill, Clark, and Stark 1975; Harris and Wolpert 2006). Thus, saccades are critical for vision, but they also produce challenges for vision, such as the need to suppress visual blur and integrate visual information across fixations (Bell et al., 2005; Melcher & Colby, 2008).) Thus, saccades and vision interact at various cognitive levels (Burr & Morrone, 2011; Eagleman, 2005; Morrone, Ross, & Burr, 2005; Ross et al., 2001). Consistent with this, saccade-related modulations are found throughout cortex (Barash et al., 1991; Berman et al., 1999; Chen & Crawford, 2017; Fairhall et al., 2017; Medendorp et al., 2007; Paré & Wurtz, 2001). But despite numerous studies on this topic in both animals and humans (summarized below), the motor and sensory influence of saccades on cortical network dynamics remains untested at the whole brain level.

Cortical saccade modulations can be roughly categorized into two categories, the first being their motor function. The cortical structures for saccade generation were first described in animals, including the parietal eye fields (lateral intraparietal sulcus; LIP) and the frontal/supplementary eye fields (FEF/SEF)), located in the lateral and medial prefrontal cortex respectively (Anderson et al., 1994; Barash et al., 1991; Hanes, Thompson, & Schall, 1995; Schall & Hanes, 1993). When a delay is interposed between visual stimulus appearance and the saccade (as in many behavioral paradigms) these areas typically show a visual response, followed by memory delay signals, and then ‘build up’ activity leading up to the saccade motor response (Powell & Goldberg, 2000; Sajad et al., 2016). These areas are also heavily interconnected with subcortical structures such as the superior colliculus, which shows a similar sequence of responses (Sadeh et al. 2015, 2018). Functional magnetic resonance imaging (fMRI) studies have confirmed the existence of human analogs to the frontal and parietal eye fields (Anderson et al., 1994; Berman et al., 1999; Levy et al., 2007; Vesia & Crawford, 2012).

The second type of saccade-related modulation in cortex relates to sensory processing. The motor signals described above are also thought to influence sensory processing, through corollary discharge (also known as ‘efference copies’). These signals are thought to suppress visual processes around the time of saccades (Berman and Wurtz 2010; Cavanaugh et al. 2016)(Bremmer et al., 2009; Burr, Morrone, & Ross, 1994; Frost & Niemeier, 2015; Kovalenko & Busch, 2016; Shioiri & Cavanagh, 1989; Thiele et al., 2002), and second, to update or “remap” presaccadic signals so that they correctly align with post-saccadic visual input (Burr & Morrone, 2011; Duhamel, Colby, & Goldberg, 1992; Melcher, 2007; Sapir et al., 2004). Such signals (directly, or through their influence on vision) have been observed throughout occipital, parietal and frontal cortex, just before and during saccades (Sommer & Wurtz, 2002; Duhamel et al., 1992; Nakamura et al. 2002; Kusunoki & Goldberg, 2003). Analagous saccade-dependent visual modulations of object location have also been observed in human fMRI studies (Merriam et al. 2007; Medendorp et al. 2003, 2005).

Finally, it is thought that the brain has specific neural mechanisms for integration of pre and post-saccadic object features (Burr & Morrone, 2011; Fabius et al,, 2020; Melcher, 2007; Prime et al. 2008, 2010). These remapping, updating and integration signals are thought to be related to perceptual distortions (especially perception of time and space) that occur before, after, or across saccades (Binda et al., 2009; Burr et al., 2011; Eagleman, 2005; Morrone et al., 2005; Ross et al., 2001); However, a recent experiment (which included behavioral data derived during the current neuroimaging study), showed that these saccade-related effects can interact with the influence of other visual stimuli. In particular, saccadic time distortion was canceled when a series of repeated stimuli were presented before saccades (Ghaderi, Niemeier, & Crawford, 2021). This finding suggests that the neural mechanisms involved in perisaccadic perception are influenced by the need to simultaneously process repeated sensory information.

Repetition of a stimulus is known to cause behavioural and neural priming effects. Behaviorally, priming effects are indexed as enhanced accuracy of recognition and faster reaction time to repeated stimulus (Horner and Henson 2008). Furthermore, duration of a repeated stimulus is perceived to be shorter than the duration of the first stimulus in the series (Eagleman and Pariyadath 2009). Neurophysiological studies showed that these behavioural effects are accompanied by modulations of neural activity in widely distributed brain areas (Summerfield et al. 2008). The neural suppression associated with stimulus repetition (repetition suppression) has been observed in medial temporal, lateral frontal, and ventral occipitotemporal cortices (Kim 2017) but these patterns can be inverted (repetition enhancement) by task conditions such as difficulty, number of repetitions, and physical stimulus properties (Kim 2017).

Notably, the cortical regions involved in repeated information processing overlap widely with the regions that are involved in sensorimotor processing for saccades(Van Pelt et al., 2010). Consistent with this, some occipital and parietal areas, particularly the cuneus and the right supramarginal gyrus, show saccade-dependent interactions between pre- and post-saccadic stimulus features akin to repetition suppression / enhancement (Baltaretu et al., 2020; Dunkley, Baltaretu, & Crawford, 2016). Therefore, these two phenomena (saccade related and repetition related distortions in visual information processing) may interact through widely distributed functional brain networks.

The neural mechanisms involved in saccade and repeated information processing, and their interaction, have thus been investigated on local, regional, and behavioral levels, but the network topology and dynamics of these phenomena, and their interactions, have not been studied at the whole-cortex level. Neurophysiology has high spatial and temporal resolution but is usually limited to local recordings. fMRI has whole-cortex capacity and reasonable spatial resolution, but it does not provide analysis in separate frequency domains and its temporal resolution is too limited to isolate presaccadic vs. post-saccadic effects. In particular, fMRI does not provide enough data points within the brief time span before saccades to do the correlations required for functional connectivity analysis. In contrast, magnetoencephalography (MEG) and electroencephalography (EEG) provide frequency band analysis and have high temporal resolution, and thanks to recent improvements in EEG source localization they now attain a spatial resolution that comes close to standard univariate fMRI analysis(Asadzadeh et al., 2020; Mégevand & Seeck, 2018; Wang et al., 2021).

Previous EEG/MEG studies have shown that the phase and amplitude of different frequency bands may be altered in the perisaccadic interval or in the face of repeated series of stimulus (Baldeweg 2007; Garrido et al. 2009) (Busch, Dubois, & VanRullen, 2009; Fabius et al., 2020; Kovalenko & Busch, 2016; McLelland, Lavergne, & VanRullen, 2016; Medendorp et al., 2007; Staudigl et al., 2017), but did not study topology and dynamics of cortical networks constructed from correlated, source localized EEG signals.

Another technical barrier is the need for analytic tools that can quantify network dynamics at the whole brain level. Recently, various approaches in graph theoretical analysis (GTA) have been proposed in neuroimaging studies to investigate various aspects of functional brain networks (Bassett & Sporns, 2017; Ghaderi et al., 2020; Rubinov & Sporns, 2010; Sporns, 2014) (Note that functional connectivity refers to signal sharing / processing, it does not imply direct anatomic connectivity). In GTA, whole brain signal correlation patterns can be treated as functional networks composed of nodes (brain regions) and edges (a functional connectivity measure between pairs of regions). GTA can be used to derive the location of network hubs (nodes with particularly high levels of connectivity) and various quantitative measures of functional network topology and dynamics (Bullmore & Sporns, 2009; Ghaderi, et al., 2020; Rubinov & Sporns, 2010; Stam, 2014), such as *clustering*, *efficiency, centrality, and energy* (see Table 1). These measures provide important quantitative information about functional network dynamics and topology, such as local vs. global connectivity, network stability, and uniformity of network connections (Bassett & Sporns, 2017; Ghaderi et al., 2020). To our knowledge, this approach has not been applied to the cortical saccade system, or its interactions with ambient visual signals.

Here, we combined EEG and GTA to study how sensorimotor signals (saccades, stimulus repetition and their interactions) influence cortical network topology and dynamics in the presaccadic period. To test the influence of stimulus repetition, we employed a classic stimulus repetition paradigm (Grill-Spector, Henson, & Martin, 2006; Verfaellie et al., 2004) where visual stimuli were repeated, followed by the same or a novel stimulus just before a saccade (Dunkley et al. 2016, Baltaretu et al. 2020). To probe interactions between presaccadic motor and visual signals, we employed the well-known effect of stimulus repetition on perception of a subsequent novel stimulus (Grill-Spector et al. 2006), by inserting this into the presaccadic interval (Ghaderi, Niemeier, & Crawford, 2021). Based on previous EEG/MEG studies (Jerbi et al. 2009; Kovalenko and Busch 2016; McLelland et al. 2016), we analyzed EEG sources in various frequency bands (i.e., theta, alpha, beta, and gamma bands). Then, we computed a functional connectivity measure (*lagged coherence* between cortical nodes) to construct functional networks.

This sequence of analyses allowed us to 1) compare cortical regions derived from source location versus network hubs derived from GTA, both with each other and the published neuroimaging literature, and 2) compute various network properties, such as topology, complexity, and stability of synchrony that were computed for these networks. We hypothesized that 1) saccades would alter local current density (related to neural activity) in the parieto-frontal regions discussed above, 2) these effects would interact with the influence of stimulus repetition on a pre-saccadic visual stimulus (Ghaderi, Niemeier, & Crawford, 2021), and that 3) at the network level, saccades would increase synchronization and integration within a functional brain network that includes parieto-frontal hubs, but engages a much broader influence on cortical visual processing.

## 2 Method

### 2-1 Participants and ethics approval

Twenty-one volunteers (12 females, aged between 18 and 44) participated in this study. All participants were right-handed and had normal or corrected to normal vision. We asked participants to report if they had a history of neuropsychiatric disorders or seizures and volunteers with positive history were excluded from the study. Each participant was paid 15 CAD/hour for participation. All participants signed an informed consent form that described all procedures about the experiment, data collecting/storage and analysis. This study was approved by the office of research ethics (ORE) at York University and the University of Toronto. All the methods in this study were performed in accordance with declaration of Helsinki.

### 2-2 Procedures and setup

A *saccade* and a *fixation* task were presented both during two separate sessions. In each session, participants were asked to perform both tasks each day, and the order of tasks was counterbalanced across participants. For each participant, the second session was conducted one week after the first. During each session participants were seated in a dark room on a comfortable chair while we recorded their EEG and electrooculography (EOG) signals (controlled online by one author, A. G., from a separate room). Thirty-five centimetres in front of them a LED monitor (25”, refresh rate 144 Hz, full HD) presented stimuli programmed in MATLAB (ver. R2019a) using the Psychtoolbox-3 (version 3.0.16)(Kleiner 2010; Kleiner, Brainard, and Pelli 2007).

### 2-3 Stimuli and trials

Saccade and fixation trials were administered in different blocks. In saccade blocks (Fig. 1-a, left; also see Ghaderi, Niemeier, & Crawford, 2021), trials started with the participant fixating a white cross (2 deg across) in the centre of a black screen and then pressing the space bar. After 300ms, a reference stimulus appeared for 30, 50, 90, or 110 ms with its upper edge 5 deg (and its centre 11 deg) below the fixation point. The reference stimulus consisted of three white parallel lines (2° wide), separated by two 3° spaces, total size 12° ×12°) that were either all horizontal or vertical. For a third of the trials the stimulus appeared only once. Another two thirds of trials had the stimulus reappear for a second and third time, respectively, with a temporal gap of 300 ms between appearances. Two hundred milliseconds after the last presentation of the reference stimulus the fixation cross jumped 10 deg randomly to the left or to the right (the total number of saccades to the left and saccades to the right was equal). After 100 ms, and before the participant was able to move their eyes, a test stimulus appeared for 70 ms (Figure 1-b). The test stimulus was identical to the reference stimulus, except the lines were rotated by 90 deg (i.e., they were vertical when the reference lines were horizontal and vice versa). After the saccade, a blue square was presented at the center of screen and participants dragged the square with a mouse to judge whether duration of the test stimulus was longer (dragging up) or shorter (dragging down) than the last reference stimulus. The configuration of the tasks in this study was similar to our previous psychophysics and modeling study that was performed with longer intervals and vertical saccades (Ghaderi, Niemeier, & Crawford, 2021). In total 720 saccade trials were collected across the two sessions. Another 720 fixation trials were collected where stimuli and procedures were the same except the fixation cross always remained in the centre of the screen (Fig. 1-a, right side).

**Figure 1:**
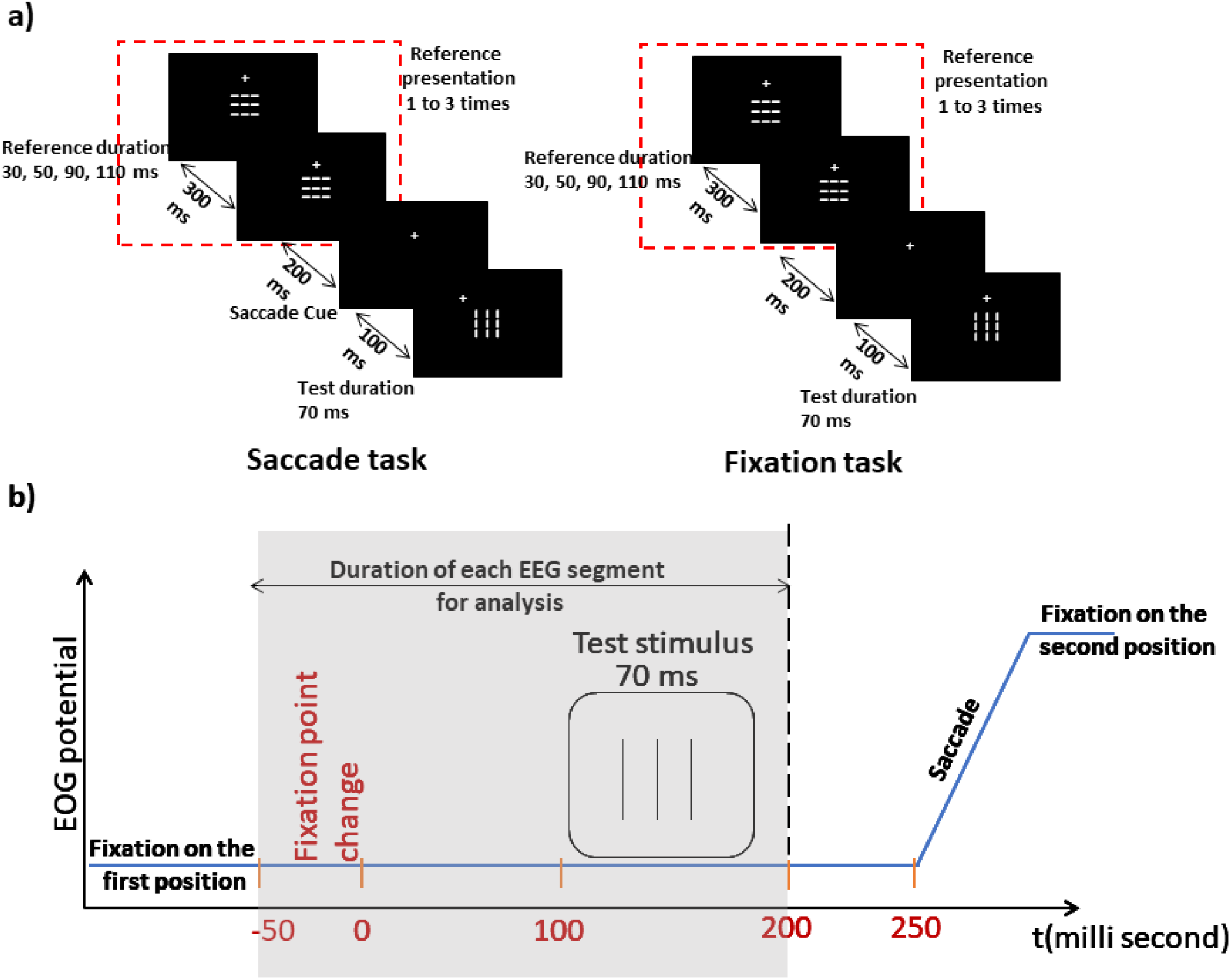
Task configurations. a) the reference stimulus (with varied duration) was presented 1 to 3 times, then the test stimulus (with fixed duration) was shown. The orientation of the reference and test stimulus were always opposite. The interstimulus interval was 300 ms. In the *saccade* task, participants were cued to perform a saccade 100 ms before the presentation of the *test* stimulus. The *fixation task* was the same overall, but participants continued fixating after stimulus presentation. b) A schematic representation of the timing of events and EEG analysis in the saccade task. EEG signals were analyzed for 200 ms across the test stimulus period, which fell in the presaccadic interval of the saccade task (or equivalent period for fixation task).

### 2-4 Saccade measurement and exclusion criteria

A schematic of the time course of saccades relative to other events is shown in Figure 1-b. EOG signals were recorded at 2048 Hz using a BioSemi System. Horizontal eye movements were monitored with bipolar EOG electrodes to the left and right sides of right eye. Vertical eye movements and blinks were detected using another electrode pair above and the eye. To distinguish microsaccades or drift from acceptable saccades, a calibration session for saccade amplitude was performed before the main experiment. In this session participants were asked to execute several 10° saccades to the left or right side (by changing the location of a fixation point on the screen). The mean change of amplitude in the EOG was calculated for 20 of these saccades. 70% of this mean was used as the minimum amplitude for saccades in the experimental data. Two other criteria for saccade exclusion were duration and latency (the time between shifting the location of fixation point and saccade onset). Saccade onsets and offsets were determined using an algorithm implemented in MATLAB code. This algorithm also excluded trials that included 1) saccade latencies outside of a 200-450 ms window, 2) saccades that took longer than 100ms, 3) multiple saccades and/or 4) blinks.

### 2-5 EEG acquisition, preprocessing, and extraction of event related signals

EEG acquisition was performed at 2048 Hz with a BioSemi system (ActiveTwo: DC amplifier, 24-bit resolution) via a 64 EEG montage. All electrodes were referenced to a Common Mode Sense (CMS) active electrode and a Driven Right Leg (DRL) passive electrode (see http://www.biosemi.com). During the recording sessions, the impedance between electrodes and scalp was kept under 5 kilohms. An online anti-aliasing first order analog filter and an online fift^th^ order lowpass filter (<134 Hz) was applied by the BioSemi amplifier. The acquisition program was the Actiview program that is an open-source program written in LabVIEW (The National Instruments Inc.). After recording, EEGs were imported to the MATLAB 2019a by EEGLAB toolbox (https://sccn.ucsd.edu/eeglab/index.php) and this toolbox was used for EEG preprocessing. A fift^h^ order bandpass digital filter (0.1 to 40 Hz) was applied.

Then, several additional steps were taken to ‘clean up’ the EEG data and remove/correct artifacts. First, to remove EEG signals with continuous artifacts, the channel statistics approach in EEGLAB (http://eeglab.org/tutorials/06_RejectArtifacts/cleanrawdata.html) was used and single channels with continuous artifacts were excluded (see section 2-6 for extracted channels). In this approach, flat channels, channels with large amounts of high frequency noise (based on their standard deviation), and channels that exhibited very low linear correlation with nearby channels (i.e., that may be caused by blinks, loose electrodes, etc.) were removed. After that, according to the criteria described in section 2-4, EOG data was inspected and trials with eye-movements / blink artifacts were automatically excluded (in MATLAB 2019b). EEG signals were then visually inspected and trials with any remaining obvious biological or technical artifact were excluded from the analysis. Finally, after excluding these trials and channels, we performed an independent component analysis (ICA) to remove any remaining muscle activity and eye movement/blink component from signals. In this step, we removed maximum four components with highest amplitude over ears (that was related to frontalis and temporalis muscle activity artifact) and eyes (that was related to eye blink/movements)(Sanei and Chambers 2013).

After removing artifacts, EEG segments were extracted. Two types of EEG segments were extracted for analysis: a 250ms duration segment *before* the onset of the first reference stimulus (used as a baseline) and another 250ms segment that contains 200ms started *after* the saccade cue and 50ms *before* the saccade cue (with 250ms signal length then the frequency resolution is 4Hz). Since, after our exclusion criteria, the saccade always started 200-450ms (mean 263.7ms ±41.4ms SD) after the cue, the latter data segment always fell within the presaccadic interval, and could never contain artifacts related to the saccadic eye movements (Jerbi et al., 2009; Keren, Yuval-Greenberg, & Deouell, 2010). A part of this presaccadic data segment aligned with the duration of the test stimulus, i.e., 100-200ms after the saccade cue, when the influence of previous stimulus repetition on sensory processing should occur (see Figure 1B for timing). To compare these with fixation condition, we selected the corresponding 250ms time periods from the fixation data, i.e., a baseline 250ms before the first reference stimulus onset, and a second segment (corresponding to the presaccadic segment) from 150ms before the test stimulus to the 100ms after test stimulus onset.

### 2-6 Excluded participants and channels

Participants with more than 60% false trials (according to both EEG and saccade criteria) were excluded from the study. In the remaining participants, there was no significant difference in percentage of excluded trials between the *saccade* task (mean: 32.3, SD=10.8) and the *fixation* task (mean= 21.9, SD=7.1). Some EEG channels had to be eliminated (because of continuous artifact, bad channel, or dead channel) in some participants. These were channel PO4 for participants #8, #13, #14, #18, channel P9 for participant #3, #14, channel TP7 for participant #17, and channel CP6 for participant #11. The excluded channels were eliminated from both tasks to equalize number of channels among conditions.

### 2-7 sLORETA source localization

The clean EEG segments were imported to the LORETA software (version 20190617) (http://www.uzh.ch/keyinst/loreta) and EEG source localization was performed in the frequency domain (cross-spectrum analysis) according to instantaneous discrete and linear solutions for the EEG inverse problem (Pascual-Marqui, 2002). Although LORETA does not have the spatial resolution of fMRI, recent work suggests that the accuracy of modified LORETA algorithms (e.g., standard LOERTA) approaches fMRI (Asadzadeh et al., 2020; Mégevand & Seeck, 2018; Wang et al., 2021). Based on the high temporal resolution of EEG, LORETA algorithms can reveal neural oscillatory activity in a wide range of frequencies including higher frequency bands, i.e., beta and gamma.

LORETA computes EEG current source density in 2394 gray matter cortical voxels. In this algorithm, distribution of EEG current source density is estimated according to a smoothness approach. This smoothness algorithm inspired the neurophysiological fact that neighboring neuronal populations exhibits local synchrony and correlated activity (Pascual-Marqui et al., 1999). The estimated current source density in voxels then are grouped in 84 ROIs, based on the Talairach coordinates of Brodmann areas (BA) obtained from a previous study (Pascual-Marqui 2002). Grouping the voxels within Brodmann areas increased statistical power while simultaneous avoiding overestimating spatial resolution to specific voxels. Thatcher et al., used LORETA and showed that source localized EEG functional connectivity between these ROIs (Brodmann areas) is correlated with synaptic density (as obtained by diffusion spectral imaging) between them (Thatcher, North, and Biver 2012). Furthermore, since we performed network analysis and we calculated functional connectivity between all pairs of regions (described in section 2-8), grouping of voxels helps to avoid spurious correlations between very close neighbour voxels.

In this study we selected all Brodmann areas that were already matched with the Talairach atlas in previous investigations (Pascual-Marqui et al. 1999; Thatcher et al. 2012). These regions contain BAs 1-11, 13, 17-25 and 27-47 in both hemispheres. Sources for the theta (4-8 Hz), alpha (8-12 Hz), beta (12-28 Hz), and gamma (28-40 Hz) bands were calculated by the standard LORETA (sLORETA) algorithm (Pascual-Marqui, 2002). Current density in these sources was compared between conditions. After that the connectivity between sources was calculated for graph theory analysis.

### 2-8 Lagged coherence and adjacency matrix

To quantify functional connectivity between brain regions, we investigated oscillatory activity of current densities in different cortical regions using sLORETA-based *intracortical lagged coherence*. *Intracranial lagged coherence* between all pairs of EEG sources (localized based on the Brodmann areas described in section 2-7) was computed in different frequency bands in sLORETA (Pascual-Marqui, 2007). According to this procedure, *lagged coherence* between two multivariate timeseries (here, oscillations of current densities in two cortical regions) in a specific frequency band (*i.e.*, *X_ω_*, *Y_ω_*; *where ω is frequency*) is defined as:

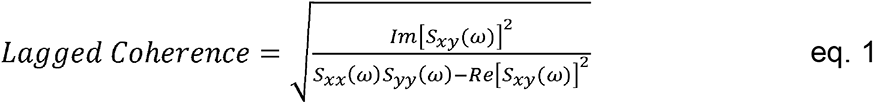

Where, *Im* and *Re* stand for the imaginary and real parts of Fourier transforms, respectively. *S_xy_*(*ω*) is cross-spectral density between *X*ω and *Yω*, and *S_xx_*(*ω*) and *S_yy_*(ω) are auto-spectral density for *X_ω_* and *Y_ω_*, respectively(Pascual-Marqui, 2007; Pascual-Marqui et al., 2011).

*Intracortical lagged coherence* is a modified version of instantaneous coherence that removes effects of volume conduction (Ghaderi, et al, 2018a; Pascual-Marqui et al., 2011). Volume conduction in EEG is generated by instantaneous changes in local field potentials and is related to electromagnetic induction in the brain tissue instead of connectivity via neural circuits (Nunez et al. 1997). The effects of volume conduction are thought to be associated with the real part of coherence in Fourier transform (Pascual-Marqui et al., 2011). In calculation of *lagged coherence,* imaginary part of Fourier transform is considered, and the real part is removed (eq. 1). In this way, the effect of volume conduction in connectivity between regions is minimized.

### 2-9 Functional brain networks: topological indices, energy, and entropy

In most brain network studies, brain regions are considered as nodes and the structural or functional relations between brain regions are treated as edges. Studies that describe physical pathways (i.e., white matter) between regions deal with *structural brain networks* and studies that only consider the linear/nonlinear relationships between neural activity in different regions describe *functional brain networks* (Rubinov and Sporns 2010). In a *weighted network*, the values of these relationships vary continuously. Here, we investigated weighted functional brain networks in several conditions (e.g., saccades vs. fixation, stimulus repetition). To model these networks, each brain region (84 Brodmann areas), was treated as a node and the *intracortical lagged coherences* (as a measure of functional connectivity) between pairs of node were treated as *edges* (Bullmore & Sporns, 2009; Ghaderi et al., 2018a; Jouzizadeh, et al., 2021). Lagged coherence was computed for each participant, condition, and frequency band. These values were then arranged in an *adjacency matrix*. In the adjacency matrix, each row / column represents one region, and the intersection of rows and columns represents the value of the *lagged coherence* between these two regions. We normalized these values to obtain consistent (weight conserving) GTA measures across conditions (Rubinov and Sporns 2011). This ensures that GTA indices are independent of the distribution of matrix weights. These adjacency matrices were used for further analysis using conventional and spectral graph theoretical approaches to obtain topological and dynamical features of functional brain networks.

#### 2-9-1 Topological features of functional brain networks

Conventional GTA indices were used to describe the topology of functional brain networks (Bullmore and Sporns 2009; Rubinov and Sporns 2010). Topological features in a complex network are characterised by the architecture of the paths between nodes, considering all nodes and edges. We used the three topological features that are most commonly used to describe human brain networks: *segregation*, *integration,* and *centrality* (Bullmore & Sporns, 2009; Rubinov & Sporns, 2010; Deco et al., 2015; Ghaderi et al., 2018b, 2019; Kitzbichler et al., 2009; Rubinov & Sporns, 2010; Sporns, 2013). These network features are described qualitatively in the following paragraphs (see Table 1 for formal mathematical definitions).

*Functional segregation* is related to the presence of highly interconnected nodes via local circuits, i.e., *modules* (Rubinov and Sporns 2010). Networks with the capacity for local / modular processing are considered to be segregated networks. Several studies have suggested that functional brain networks are segregated networks (Bassett & Bullmore, 2007; Bullmore & Sporns, 2009, 2012; Watts & Strogatz, 1998). In functional network analysis, the c*lustering coefficient (CC)* is often used to measure the level of segregation (Bialonski & Lehnertz, 2013; Deco et al., 2015; Ghaderi et al., 2018b; Pedersen et al., 2015; Rubinov & Sporns, 2010; Sjoerds et al., 2017). The clustering coefficient is calculated for each node and the average *CC* for all nodes is used to quantify the overall level of functional segregation in the network. For each node, *CC* is related to number of clusters around the node. In this study, clusters are defined as ‘triangles’, i.e., triplets of interconnected neighbour nodes. In a weighted network, when the neighbors of a node all have maximum connection weights, *CC* is maximum and equals 1, whereas *CC* equals 0 if the node has no connections.

*Functional integration* is related to combination of information across distributed network nodes, and indicates the level of global information processing (Rubinov and Sporns 2010). Several studies have shown that functional brain networks are highly integrated and exhibit very fast integration between distributed anatomical regions (Bassett & Sporns, 2017; Bullmore & Sporns, 2012; Rubinov & Sporns, 2010; Stam, 2014; Stam & Reijneveld, 2007). Functional integration in brain networks can be affected by alterations in brain states, diseases or disorders (Ghaderi et al., 2017, 2020; Hasanzadeh et al., 2020; Rubinov & Sporns, 2010; Stam & Reijneveld, 2007; van den Heuvel et al., 2008). In GTA, functional inegration can be measured by *global efficiency* (*Ef*), which is inversely proportionate to the average highest weights between all pairs of nodes (Boccaletti, et al., 2006; Newman, 2008; Rubinov & Sporns, 2010). In a network where all node pairs are highly connected, path lengths are called ‘short’ and thus information can be transferred rapidly. Such networks have high global efficiency, i.e., high levels of integration (Boccaletti et al. 2006; Newman 2008).

Networks with high values of both *CC* (clustering) and *Ef* (integration) are known as *small world* networks. Such networks display fast and efficient information processing. Many studies have suggested that brain networks are small-world networks (Bassett & Bullmore, 2006; Ghaderi et al., 2018a; Muldoon et al., 2016; Watts & Strogatz, 1998). Small-worldness in functional brain networks is affected by both specific cognitive states (Li et al. 2019; Lin et al. 2018) and brain diseases / disorders (Jouzizadeh et al. 2021; Lynall et al. 2013; Stam 2014). *Small-world propensity (SW*) has been used to evaluate small-worldness in functional brain networks (Muldoon et al. 2016). SW is calculated by taking the ratio of *CC* and *Ef* in the experimental brain network in comparison to these measures in control networks (i.e., random and regular networks), where higher SW values indicate higher small-worldness (Muldoon et al. 2016).

Finally, *centrality of nodes* is used to evaluate the importance specific nodes. Nodes that are connected to many other nodes are called ‘important’. Nodes that have many connections to important neighbors are called hubs. Hubs can be defined mathematically (Table 1) as nodes with the highest *Eigenvector centrality* (*Eig_C_*) values (Fagerholm et al., 2015; Ghaderi et al., 2021; Joyce et al., 2010). In functional brain networks, alteration of *Eig_C_* suggests that the node’s contribution to the whole network has changed. Here, we compared *Eig_C_* of all brain regions in different conditions to find which regions were most important for different brain networks, for example during saccades versus fixation.

All these topological indices were calculated using *Brain Connectivity Toolbox* (Rubinov and Sporns 2010) (https://sites.google.com/site/bctnet/).

#### 2-9-2 Synchrony and complexity of functional brain networks

Note that in EEG data, each time series measurement takes the form of an oscillation, and the entire set of measurements can be considered an oscillatory system. For such systems, each oscillatory unit can be treated as a node and the level of similarity between oscillations across nodes as the edge. A subfield of GTA, known as *spectral graph theory*, is devoted to evaluate stability of synchronizations and robustness of oscillatory systems (Spielman 2007; Wilf 1967). In spectral graph theory, *stability of synchronization* (i.e., the level of coupling between all dynamical nodes) can be evaluated by employing linear algebra and investigating the eigenvalue spectrum of the adjacency matrix (Atay et al, 2006; Larremore et al., 2011; Spielman, 2007). The sum of all eigenvalues of the matrix is called *energy of graph (H*), and is directly related to the stability of synchronization in the system (Gutman, 2013; Li et al., 2012). Here, we used *H* to evaluated stability of synchronization in our EEG data (Ghaderi et al., 2020, 2021). Previous studies showed that when all nodes in a network show high synchronization (coupling) during a specific period, then the network exhibits a high value of *H* (Daianu et al. 2015; Ghaderi et al. 2020; Gutman and Zhou 2006). In this study, we calculated *H* using the *eig* function (MATLAB R2019) for each adjacency matrix and then compared this value across conditions and participants.

To investigate *network complexity* (the variety of connections among nodes) in our functional brain networks, Shannon entropy (*S*) of networks were compared between conditions (Ghaderi et al., 2019, 2020). The value of *S* is directly related to the number of possible states for a probability system (e.g., *S* is higher for dice rolling than a coin toss). For an adjacency matrix (with many possible connectivity states between nodes), *S* is higher in the presence of more variable connection weights. *S*=0 when all values of connections are same, whereas it increases in presence of rare values, called ‘surprise states’ (Ghaderi et al., 2019, 2020, 2021). In a functional brain network, surprise states occur for brain functions that require high coupling between two specific nodes (Ghaderi et al., 2019, 2020, 2021). In this study we used the *entropy* function (MATLABR2019) to compute *S* for all adjacency matrices (again, see Table 1 for mathematical details).

### 2-10 Statistical analysis

In this study we aimed to investigate 1) the brain regions and network properties associated with the presaccadic signal processing, and 2) how these properties were modulated by repetition of a presaccadic reference stimulus, in a task where participants had to judge the duration of a subsequent perisaccadic visual stimulus. To this aim, four conditions were assumed. 1) The *saccade* condition i.e., trials in the *saccade* task that were presented with 0 repetitions of the reference stimulus. 2) The *fixation* condition i.e., trials in the *fixation* task that were presented without repetition of the reference stimulus. 3) The *saccade/repetition* condition i.e., trials in the *saccade* task that were presented repetitions of the reference stimulus, combined for statistical power because they did not produce different effects in a preliminary psychophysical study (Ghaderi, Niemeier, & Crawford, 2021) 4) The *fixation/repetition* condition i.e., trials in the *fixation* task that were presented with one or two repetitions of *reference*.

To compare brain activity among the different conditions, a *cluster-based nonparametric permutation t-test*(Maris and Oostenveld 2007) was performed to find significant differences of current densities within Brodmann areas. For each *cluster-based nonparametric permutation t-test,* 10000 random shuffles were implemented. The network indices were compared using the separate *nonparametric permutation t-test* (10000 random shuffles and significance level<0.05) among conditions. The sources of activity and network indices of these pairs of conditions were compared: *saccade* vs. *fixation, saccade/repetition* vs. *saccade,* and *fixation/repetition* vs. *fixation*.

Then, to correct the induced error of multiple comparisons (error type 1), false discovery rate (FDR) analysis was performed and corrected p-values under 0.05 were considered significant. The latter statistical corrections were done separately for each EEG band to compensate for three task comparisons (*saccade* vs. *fixation, saccade/repetition* vs. *saccade,* and *fixation/repetition* vs. *fixation*). The nonparametric cluster-based permutation t-test and FDR analysis were performed using MATLAB 2019a.

## 3 Results

An analysis of the behaviour and perceptual responses recorded in this experiment was presented in a separate manuscript (Ghaderi, Niemeier, & Crawford, 2021). This analysis showed that 1) saccades compressed the perceived duration of the test stimulus, 2) previous stimulus repetition dilated the perceived duration of the test stimulus and 3) the repetition enhancement effect cancelled saccadic time compression effect. Here, we analyze the corresponding brain signals in terms of the objective sensorimotor events, i.e., stimulus repetition and saccade motor signals, rather than subjective perception.

To evaluate neural processes associated with saccade preparation, EEG data from the presaccadic interval of the *saccade task* were compared to data from the same time segment in the *fixation* task (see method section and Figure 1 for more details). First, we tested the possible baseline differences between EEG signals in the *fixation* and *saccade* tasks. The results showed that there is no difference in local current density activities as well as network features between the *fixation* and *saccade* tasks. Therefore, any differences between the presaccadic interval and the corresponding fixation data (relative to their baselines) can be ascribed to saccade or saccade-repetition interaction effects.

After baseline comparisons, we compared the presaccadic interval between the *saccade* and *fixation* tasks. To test how these signals interact with presaccadic visual processing, this time interval incorporated the presentation of a test stimulus and was preceded by a single or repeated presentation of a reference stimulus (Figure 1). Thus, we considered the presaccadic interval under four conditions: *Fixation (F), Saccade (S),* both after a single presentation of the reference stimulus, as well as *Fixation/Repetition (FR),* and *Saccade/Repetition (SR)* conditions. In the following two sections we begin with source localization analysis, using fairly standard neuroimaging conventions, followed by a functional network analysis based on GTA. Each of these sections is broken down into four subsections: saccade effects, stimulus repetition effects, their interactions and then post-hoc analysis. Table 2 provides the functional anatomy and coordinates of Brodmann areas mentioned in the text.

**Table 1:**
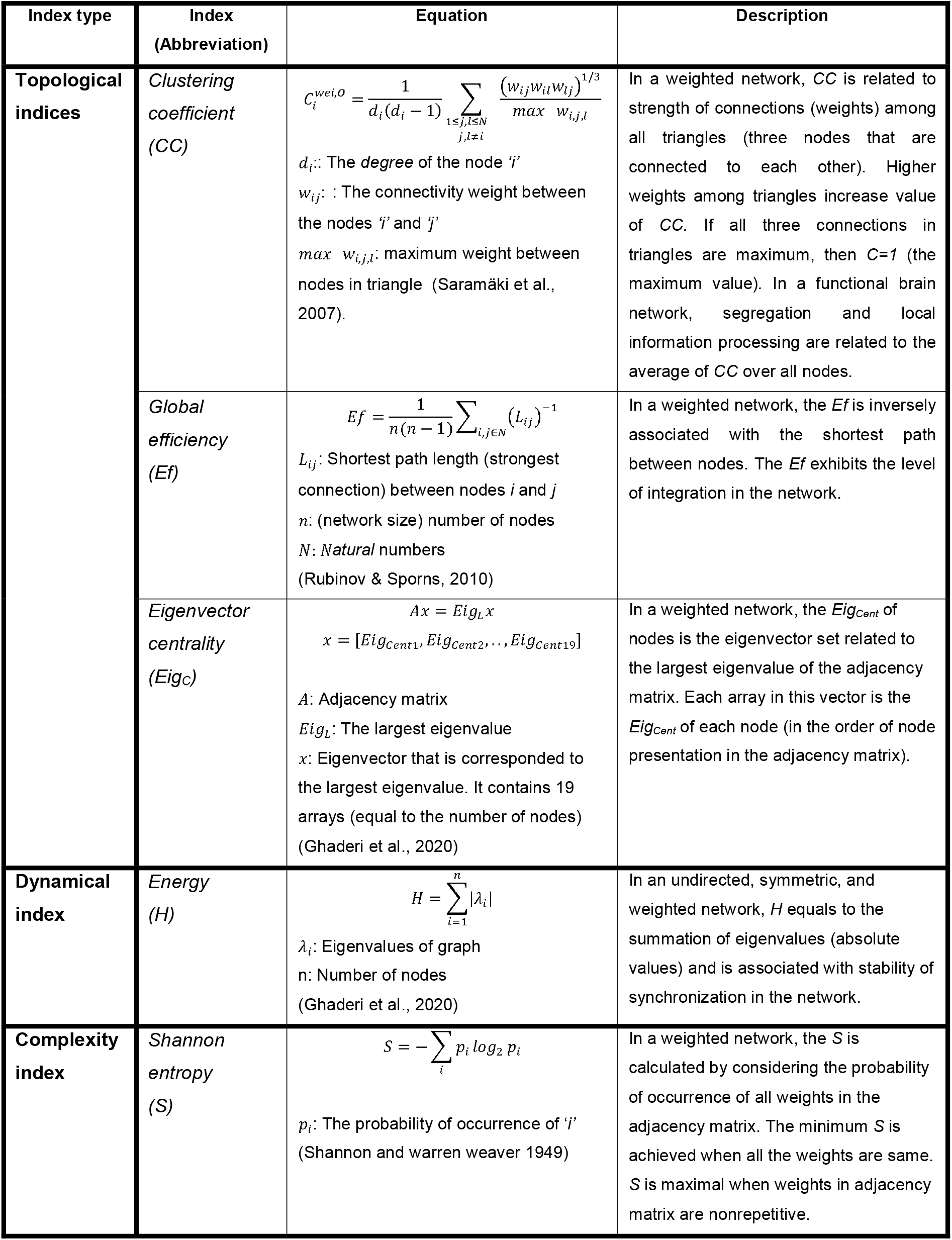
Description of graph theory indices in this study

**Table 2:** Brodmann areas and corresponded anatomical regions

| Brodmann area | Anatomical region | Talairach coordinates (X, Y, Z) |
| --- | --- | --- |
| BA 3 | BA 3 is in the postcentral gyrus and contains some regions of primary somatosensory cortex (Strotzer 2009). | (46, -19, 52) |
| BA 5, 7 | In human cortex, BA 5 and 7 are associated with posterior parietal regions. BA 5 contains superior parietal lobule and BA 7 contains precuneus and parts of superior parietal lobule (Garey 1999). | BA 5: (11, -43, 60)<br>BA 7: (11, -66, 63) |
| BA 8 | In the human cortex, BA 8 is a part of frontal cortex and contains frontal eye field region (Paus 1996). | (12, 27, 53) |
| BA 9 | In the human cortex, BA 9 is in frontal cortex and is associated with dorsolateral and medial prefrontal cortex (Rajkowska and Goldman-Rakic 1995). | (14, 46, 37) |
| BA 40 | BA 40 is a part of parietal cortex and contains inferior lateral regions of parietal cortex and especially supramarginal gyrus (Rushworth, Behrens, and Johansen-Berg 2006). | (41, -39, 52) |
| BA 44, 45 | In the human cortex, BA 44, 45 are parts of frontal cortex and contain regions of inferior frontal gyrus (Garey 1999). | BA 44: (52, 13, 15)<br>BA 45: (-38, 21, 6) |
| BA 46 | In the human cortex, BA 46 is in frontal cortex and contains the regions of dorsolateral prefrontal cortex (Rajkowska and Goldman-Rakic | (-33, 30, 12) |
|  | 1995). |  |

### 3-1 Source localization

#### 3-1-1 Saccade vs. fixation

As noted in the methods, we clustered our data voxels into ROIs based on Brodmann areas, in order to increase statistical power and avoid overestimating spatial resolution. False discovery rate (*FDR*) analysis indicated significant differences of clustered voxels between the *Saccade* and *Fixation* conditions, i.e. [(SR +S) – (FR + F). In all regions and all investigated frequencies, the current density was decreased in the *saccade* condition in comparison to the *fixation* condition (Figure 2). The statistical values of these differences are shown in Table 3. This analysis yielded significant differences for t-values higher than 4.321 in the theta, 5.540 in the alpha, 4.535 in the beta, and 4.405 in the gamma bands (Table 3). Current modulations tended to peak near the medial portions of superior frontoparietal cortex, especially in the theta and gamma bands. After *FDR* analysis and according to the sLORETA map, significant differences were observed in BAs 5, 7, 8, and 46 (Figure 2; Table 2). According to standard coordinates (Table 2), areas 8 and 46 correspond to the frontal eye fields and dorsolateral prefrontal cortex. A large area of saccade modulation was also observed in posterior parietal cortex (PPC) spanning the superior and medial portions of Brodmann’s areas 5 and 7, i.e., dorsal posterior parietal cortex, including the medial portion of PPC (precuneus). Overall, these source-localized regions spanned well-known sites obtained from neurophysiology and fMRI studies, including the frontal / parietal eye fields (Hanes et al., 1995; Hanes & Wurtz, 2001; Paré & Wurtz, 2001) and medial regions associated with more complex visuospatial tasks (Chen & Crawford, 2017; Chen et al., 2018).

**Figure 2:**
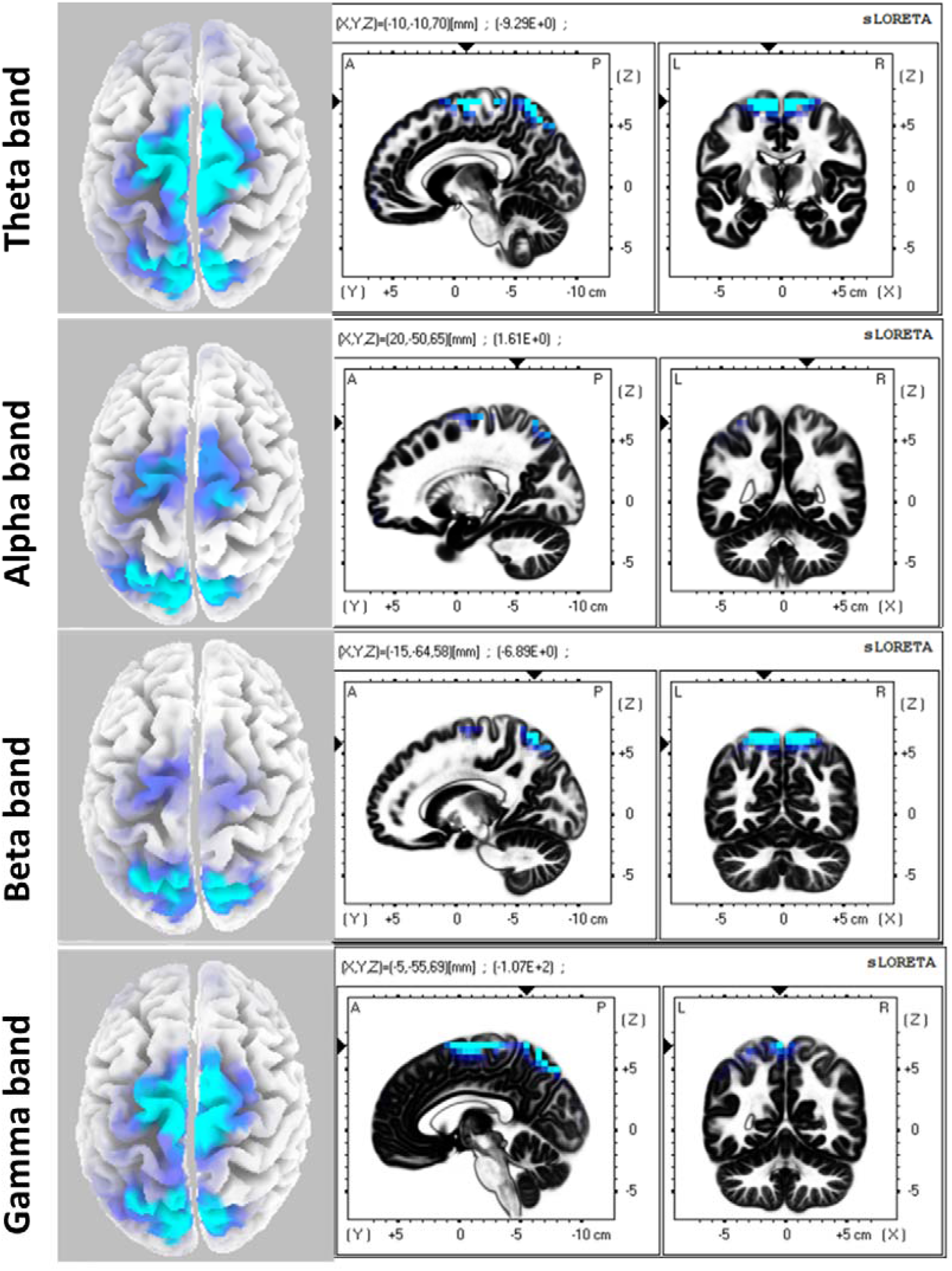
Significant EEG current density differences between the saccade condition and fixation condition for each frequency band (rows). Source localized EEG are projected (by sLORETA) onto a 3D model of the cortical surface from the above view (left column) as well as onto anatomic scans (according to sLORETA map) from the sagittal and coronal views (right columns). The blue color shows decreased current density in the *saccade* condition in comparison to the *fixation* condition. The scale of colors from dark to light blue *t-value*s from a two-tailed permutation *t-test* (lighter color shows higher *t-values*). The *t-value*s below the threshold of significant level have not been shown.

**Table 3:** The statistical details of comparisons between *saccade* and *fixation* conditions.

|  | Frequency band | Significance level | Regions with significant differences after FDR analysis |
| --- | --- | --- | --- |
| **Saccade vs. Fixation** | **Theta** | Two-Tailed (A<>B):<br>t(0.05)=4.321,<br>p=0.01040 | In the saccade condition, current density was bilaterally suppressed in superior / medial frontal, precuneus and superior parietal regions. |
|  | **Alpha** | Two-Tailed (A<>B):<br>t(0.05)=5.540,<br>p=0.00020 | In the saccade condition, current density was bilaterally suppressed in superior frontal, precuneus and superior parietal regions. |
|  | **Beta** | Two-Tailed (A<>B):<br>t(0.05)=4.535,<br>p=0.01340 | In the saccade condition, current density was bilaterally suppressed in superior frontal, precuneus and superior parietal regions. |
|  | **Gamma** | Two-Tailed (A<>B):<br>t(0.05)=4.405,<br>p=0.00020 | In the saccade condition, current density was bilaterally suppressed in superior / medial frontal, precuneus and superior parietal regions. |
| **Saccade vs. Saccade/repetition** | **Theta** | Two-Tailed (A<>B):<br>t(0.05)=4.321,<br>p=0.01040 | In the saccade/repetition condition, current density was suppressed in the superior frontal gyrus (left side) and in precuneus (both sides). |
|  | **Alpha** | Two-Tailed (A<>B):<br>t(0.05)=4.321,<br>p=0.01040 | In the saccade/repetition condition, current density was suppressed in precuneus (both sides). |
|  | **Beta** | Two-Tailed (A<>B):<br>t(0.05)=5.540,<br>p=0.00020 | In the saccade/repetition condition current density was bilaterally enhanced in the medial frontal and post central gyrus. |
|  | **Gamma** | Two-Tailed (A<>B):<br>t(0.05)=4.535,<br>p=0.01340 | In the saccade/repetition condition current density was decreased in the superior frontal gyrus (left side) and enhanced in the superior parietal lobe (both sides). |
| **Interactions: (SR-S)-(FR-F)** | **Theta** | Two-Tailed (A<>B):<br>t(0.05)=3.721,<br>p=0.01770 | The activity of SR-S was significantly higher than FR-F in bilateral precuneus region |
|  | **Alpha** | Two-Tailed (A<>B):<br>t(0.05)=4.321,<br>p=0.01040 |  |
|  | **Beta** | Two-Tailed (A<>B):<br>t(0.05)=4.535,<br>p=0.01340 |  |
|  | **Gamma** | Two-Tailed (A<>B):<br>t(0.05)=4.321,<br>p=0.01040 |  |

#### 3-1-2 Repetition vs. no repetition

Previous studies have shown that repetition of a visual stimulus alters perception of a subsequent novel test stimulus (which fell within the EEG analysis window of the current study). Here, we used this paradigm as a probe into presaccadic visual signal processing. We first tested the main influence of stimulus repetition on the combined saccade/fixation dataset, i.e., [(SR + FR) – (S + R)]. Based on nonparametric permutation t-tests, significant differences between the *repetition* condition and the *no-repetition* conditions were only observed in the beta band (t-value=-8.37, |t-value|>4.321 was significant) and were confined to clustered voxels of BA 7 (dorsal posterior parietal cortex). In this band/region, current density was lower in the repetition condition (Figure 3). We next tested interactions between this effect and saccades, to directly probe the influence of presaccadic signal processing on simultaneous visual processing signals.

**Figure 3:**
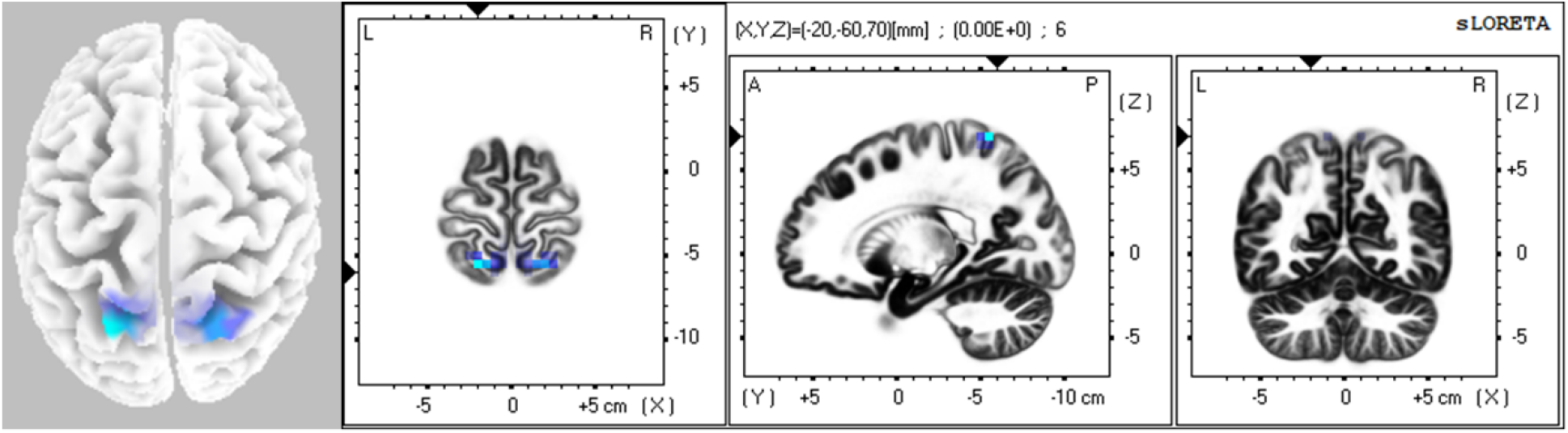
Significant EEG current density differences between the *repetition* condition and *no-repetition* for beta band. Source localized EEG are projected (by sLORETA) onto a 3D model of the cortical surface from the above view (left column) as well as onto anatomic scans (according to sLORETA map) from the sagittal and coronal views (right columns). condition. The blue color shows decreased current density in the *repetition* condition in comparison to the *no-repetition* condition. The spectrum of colors from light blue to dark blue is related to *t-value*s in two tails permutation *t-test* (lighter color shows higher *t-values*). The *t-value*s below the threshold of significant level has not been shown.

#### 3-1-3 interaction of saccade and repetition effects

To directly test the interaction between saccade and repetition effects, we compared current densities as follows: (*Saccade/repetition* minus *Saccade* condition) minus (*Fixation/repetition* minus *Fixation* condition), i.e., [(SR-S) – (FR-F)} (Figure 4). Cluster-based permutation tests and FDR analysis showed significant differences in t-values higher than: 3.721 in the theta, 4.321 in the alpha, 4.535 in the beta, and 4.321 in the gamma bands. Based on these analyses significant differences were observed in bilateral BA 7 (all frequency bands), including the superior parietal cortex and precuneus regions (Figure 4). The statistical details of these results are presented in Table 3.

**Figure 4:**
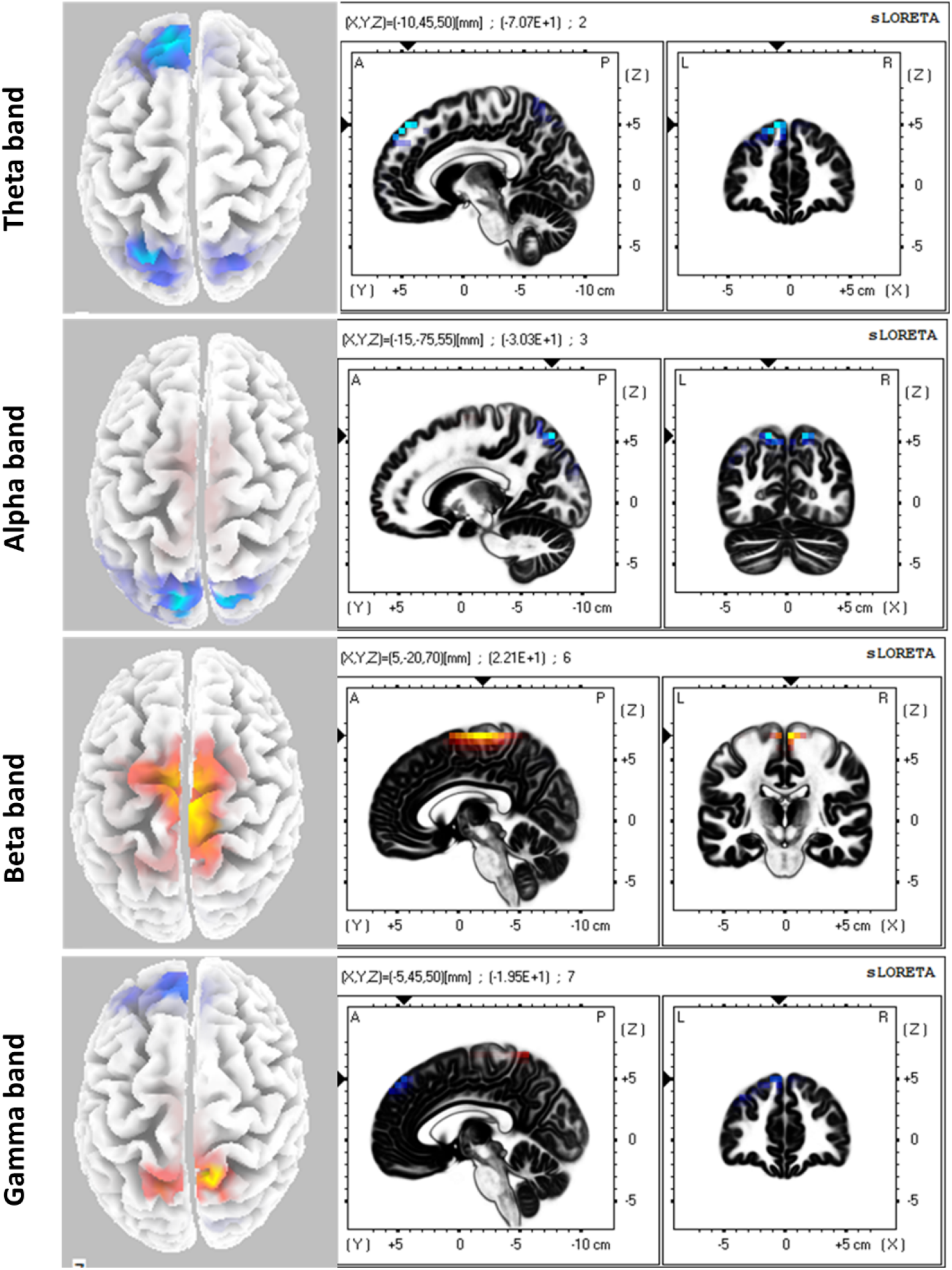
Significant EEG current density differences between the *saccade* condition and the *saccade/repetition* condition for each frequency band (rows). Source localized EEG are projected (by sLORETA) onto a 3D model of the cortical surface from the above view (left column) as well as onto anatomic scans (according to sLORETA map) from the sagittal and coronal views (right columns). The blue regions are related to decreased current density and the red and yellow regions are related to increased current density in the *saccade/repetition* condition in comparison to the *saccade* condition. The spectrum of colors from light blue to yellow is related to *t-value*s in two tails permutation *t-test* (lighter colors shows higher *t-values*). The *t-value*s below significant threshold has not been shown.

#### 3-1-4 Post hoc analysis: influence of stimulus repetition in the presaccadic interval

In a post hoc analysis we analysed FR-F and SR-S separately. First, we removed all saccade trials to compare *fixation-repetition* (FR) and *repetition* (R) trials. According to the *FDR* analysis, we observed no significant difference between trials with repetition and without repetition of reference stimulus. However, in the *saccade* task (SR – S), the *FDR* analysis showed significant differences in neural activity for clustered voxels between the *saccade/repetition* condition and the *saccade* condition (Figure 5). In this case, stimulus repetition sometimes resulted in either reduced (blue) or increased (orange) current density, depending on the region and band. The *FDR* indicated t-values higher than 4.321 in the theta, 4.321 in the alpha, 5.540 in the beta, and 4.535 in the gamma bands (Table 3). The significantly increased current density in the *saccade/repetition* condition (in comparison to the *saccade* condition) was observed in Brodmann areas 3 (beta band), and 7 (gamma band) in both hemispheres (Figure 5). BA 3 includes the postcentral gyrus and BA 7 (posterior parietal cortex) including the precuneus (Table 2). A significantly decreased current density (in the *saccade/repetition* condition compared to the *saccade* condition) was observed in left Bas 8, 46 (theta and gamma bands) and bilateral BA 7 (theta and alpha bands) (Table 3). BAs 8, 46 span superior and medial frontal gyrus including the FEF. The statistical details of these results are presented in Table 3. In short, these modulations again spanned parieto-frontal areas associated with saccades and higher level visual / executive function (Leoné et al., 2014).

**Figure 5:**
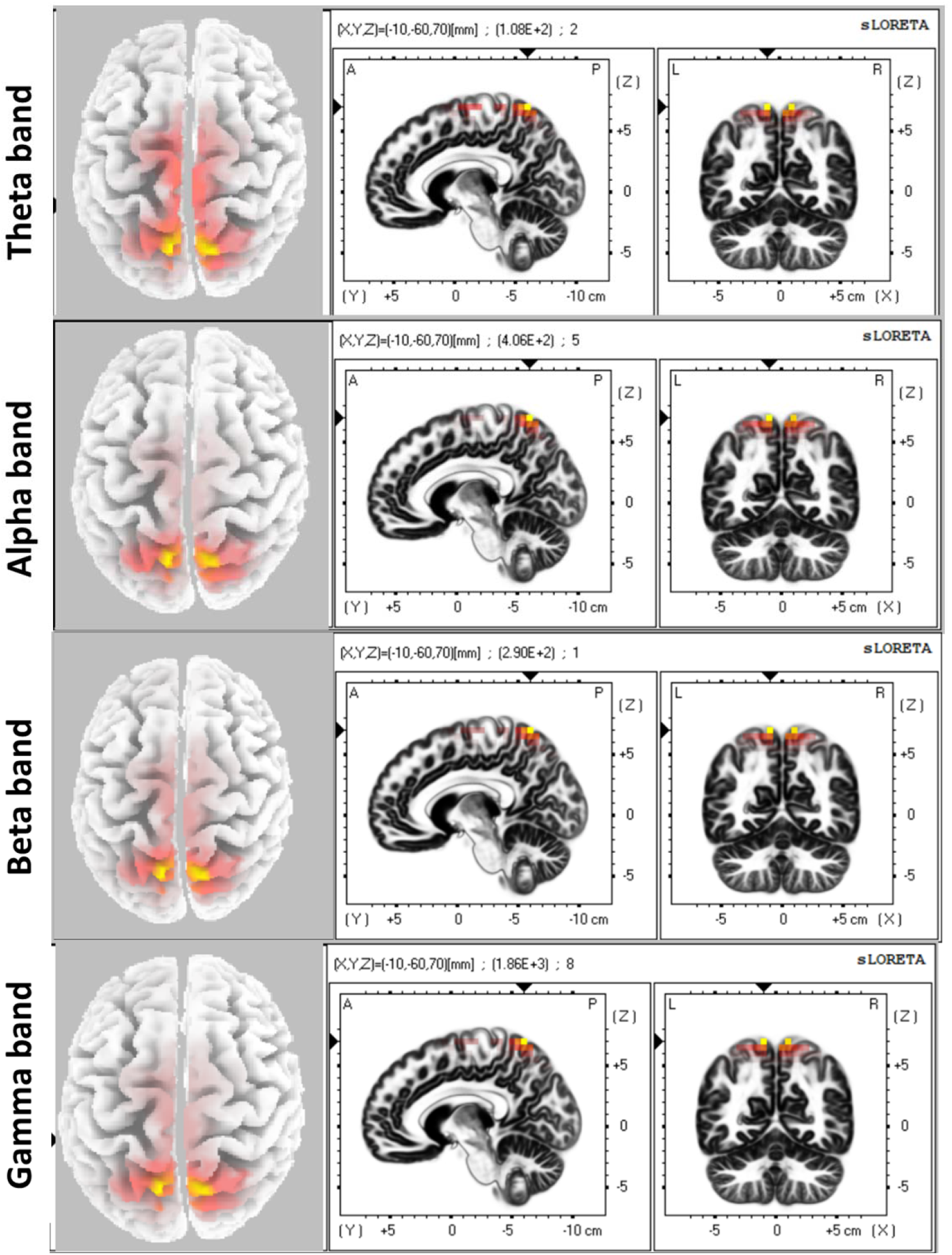
Significant EEG current density differences between SR-S and FR-F for each frequency band (rows). Source localized EEG are projected (by sLORETA) onto a 3D model of the cortical surface from the above view (left column) as well as onto anatomic scans (according to sLORETA map) from the sagittal and coronal views (right columns). The red to yellow regions are related to increased current density in the *SR-S* in comparison to the *FR-F*. The spectrum of colors from red to yellow is related to *t-value*s in two tails permutation *t-test* (yellow color shows higher *t-values* that are higher than significant threshold.

Overall, the regions of modulation observed in this study were consistent with those observed in fMRI studies of saccades (Alkan et al., 2011; Berman et al., 1999; Dunkley et al., 2016; Fairhall et al., 2017). However, source localization analysis is limited in terms of understanding the functional connectivity between these regions and with other areas of cortex in this task.

### 3-2 Network Analysis

For the next stage of our analysis, we re-examined the data using graph theoretical analysis (GTA). By this analysis we can test the architecture and dynamics of functional connectivity patterns in whole brain network in presaccadic interval. To quantify network topology and dynamics we computed the following GTA measures: 1) *eigenvector centrality (Eig_C_)* to identify important nodes and hubs, 2) *clustering coefficient (CC)* to represent the degree of segregation of the network into local clusters, *3) global efficiency (Ef)* to represent integration of signals across the network, *4) small-worldness (SW)* to represent the overall efficiency of the network at both local and global levels, 5) energy (H) to represent stability of synchronization in the network, and *5) entropy (S)* to determine the complexity of the network (see Methods sections 2.8, 2.9 for qualitative explanations and Table 1 for formal definitions of these terms). Then these values were statistically compared between conditions by permutation tests and the *FDR* approach for multiple comparisons correction. Following the same order as the source localization analysis in the previous section, we used this approach to examine functional connectivity for Saccades vs. Fixation, Stimulus Repetition vs. no Repetition, and their Interactions.

#### 3-2-1 Saccade vs. fixation

Figure 6 shows the anatomic distributions of network nodes that exhibited significant differences of *eigenvector* centrality (*Eig_C_*) in the *saccade* minus *fixation* condition, based on FDR analysis in the alpha and beta bands. In other words, this figure shows brain areas with increased functional connectivity just before a saccade. In this figure, *small black dots* indicate the sites tested and the *blue-green lines* indicate functional connections that were stronger in the saccade condition (only the top 20% are shown). The *large red dots* indicate hubs with significant differences of centrality, and the functional connections of these particular nodes are highlighted in red. In the alpha band, BA 39 (left hemisphere) (t=2.21, p=0.04) was a significant hub. BA 39 includes angular gyrus and supramarginal gyrus (SMG). This area was more widely connected to the frontocentral and occipital regions just before saccades. In the Beta band, significant differences of *Eig_c_* were observed in BA 8 (both hemispheres) (left: t=3.31, p=0.004; right: t=2.46, p=0.024) that includes FEF). These regions were more widely connected to the prefrontal and temporoparietal regions (right hemisphere) and centroparietal regions (left hemisphere). In short, this analysis revealed hubs in brain areas associated with saccade production and trans-saccadic integration), and their functional connections to occipital, parietal, and frontal cortex during this particular task (Baltaretu et al. 2020).

**Figure 6:**
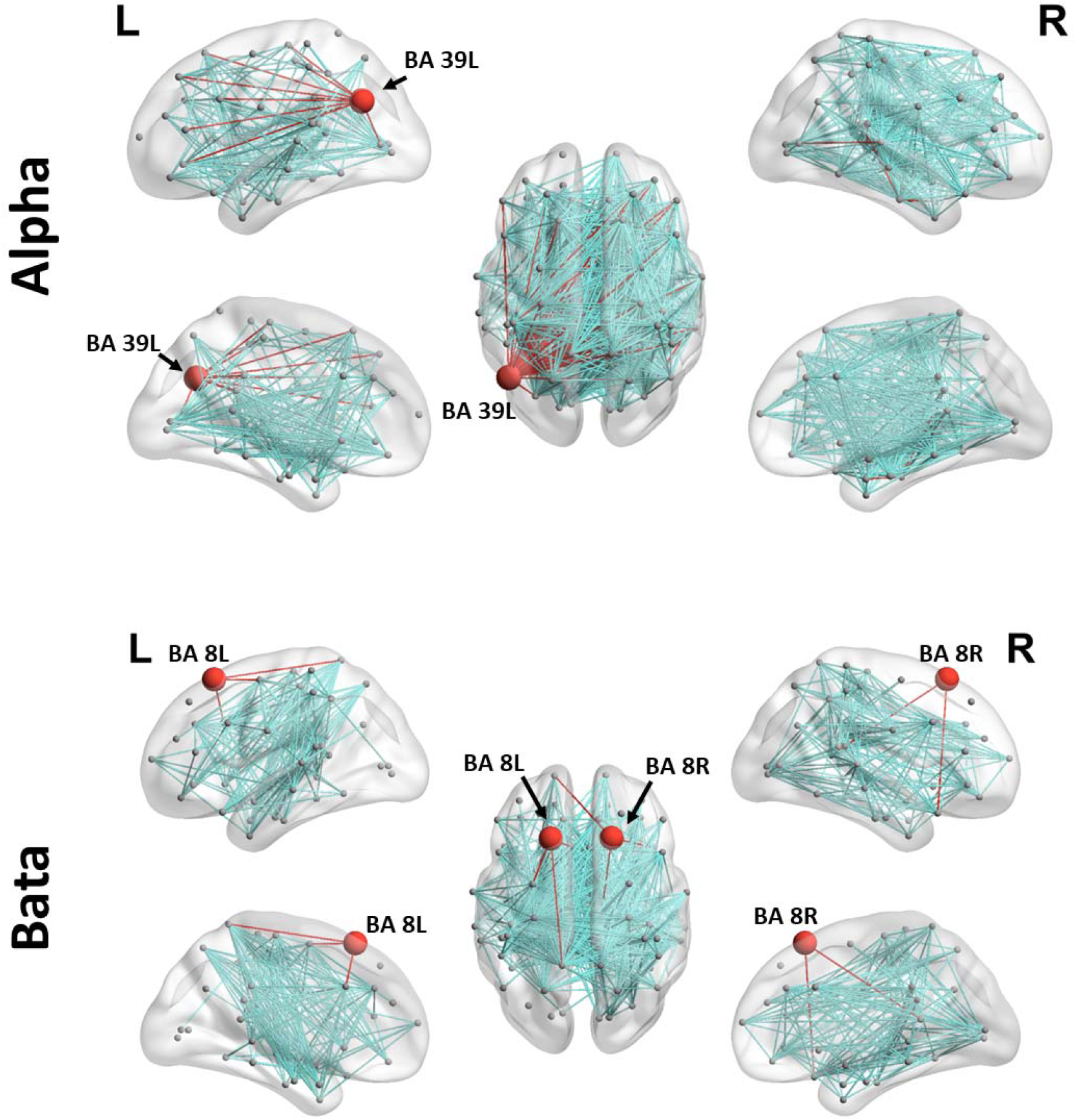
GTA connectivity maps in the *saccade* condition vs. *fixation* condition comparison. Data are projected onto lateral (upper row), upper (middle) and medial (lower row) views of the left (L) and right (R) representations of cortex (generated by BrainNet Viewer toolbox(Xia, Wang, and He 2013), MALTAB R2019). Brodmann areas that showed significant differences of centrality between conditions were considered as hubs and are shown by bolded red dots, and their connections are highlighted in red. The 20% of strongest connections related to differences between conditions (*saccade-fixation*) are shown as connectivity lines (edges), in blue-green for non-hub connections.

Whereas Figure 6 only illustrated nodes and connectivity for specific frequency bands, Figure 7 shows how five more general GTA measures changed in the saccade condition, across all bands analyzed. Each ‘violin plot’ shows the distribution of values across all participants for a given GTA measure (columns CC, Ef, SW, H, Entropy) and frequency band (rows theta, alpha, geta, gamma). Significant increases (*/**) in the clustering coefficient (CC) and in global efficiency (EF) suggest greater local and global connectivity, respectively, in presaccadic functional networks. Furthermore, the presaccadic interval was associated with a higher degree of synchronization stability and complexity, as measured by energy (H) and Shannon entropy (S). Only the measure of small-worldness (SW) remained unchanged. (See Figure Legend 7 for Statistical Values).

**Figure 7:**
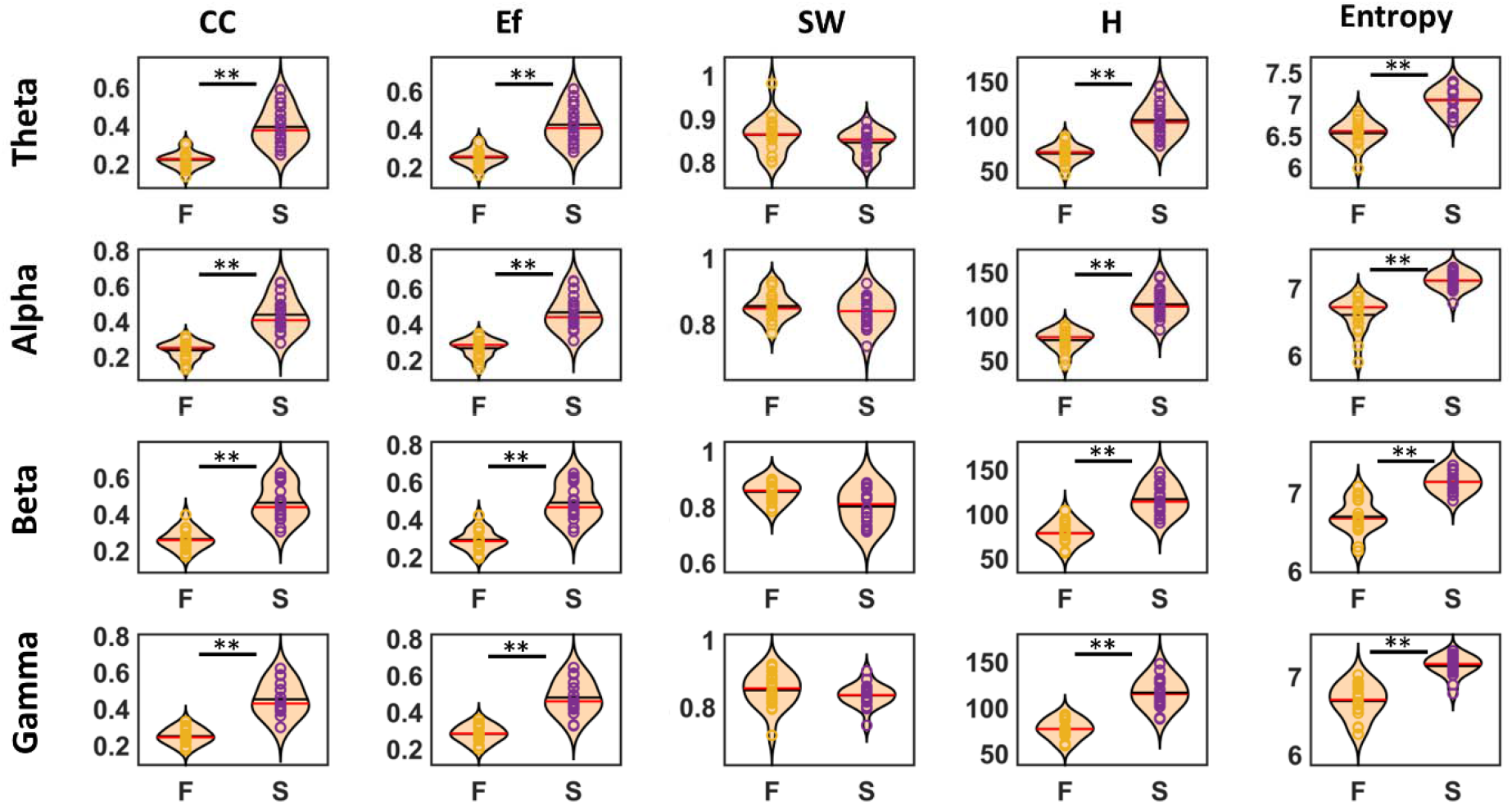
The violin bar plots of network indices in the *fixation* and the *saccade* conditions. Each dot present one individual and the width/shape of violin plots shows distributions of measures for all individuals (Violin plots combine the attributes of standard bar graphs and standard frequency histograms but arranged vertically with frequency shown bidirectionally in the horizontal dimension). The * indicates p<0.05 and the ** indicates p<0.005. F: fixation condition, S: saccade condition, CC: clustering coefficient, Ef: global efficiency, SW: smallworldness, and H: energy. Statistical details: *Clustering coefficient* (theta: t=7.18, p<0.001; alpha: t=8.78, p<0.001; beta: t=7.67, p<0.001; gamma: t=8.14, p<0.001), *global efficiency* (theta: t=7.38, p<0.001; alpha: t=9.04, p<0.001; beta: t=7.82, p<0.001; gamma: t=8.17, p<0.001), *energy* (theta: t=8.85, p<0.001; alpha: t=10.84, p<0.001; beta: t=9.32, p<0.001; gamma: t=9.36, p<0.001), and *entropy* (theta: t=10.31, p<0.001; alpha: t=9.34, p<0.001; beta: t=8.58, p<0.001; gamma: t=8.60, p<0.001).

#### 3-2-2 Stimulus Repetition vs. no repetition: main effect and interaction with saccades

The combined saccade/fixation dataset only showed one significant main effect for repetition vs. no repetition: significantly higher clustering coefficient for the no-repetition condition (t=1.86, p=0.047) in the beta band. However, several significant interactions were observed between repetition and saccade effects.

Figure 8 shows the anatomic distribution of network nodes with significant differences of *Eig_C_* between *fixation/repetition* minus *Fixation* (*FR-F*) and *saccade/repetition* minus *Saccade* (*SR-S*) according to FDR analysis in different bands. In other words, brain areas where saccade plan and stimulus repetition interacted to influence functional connectivity in the presaccadic interval. This figure follows the same conventions as Figure 6.

**Figure 8:**
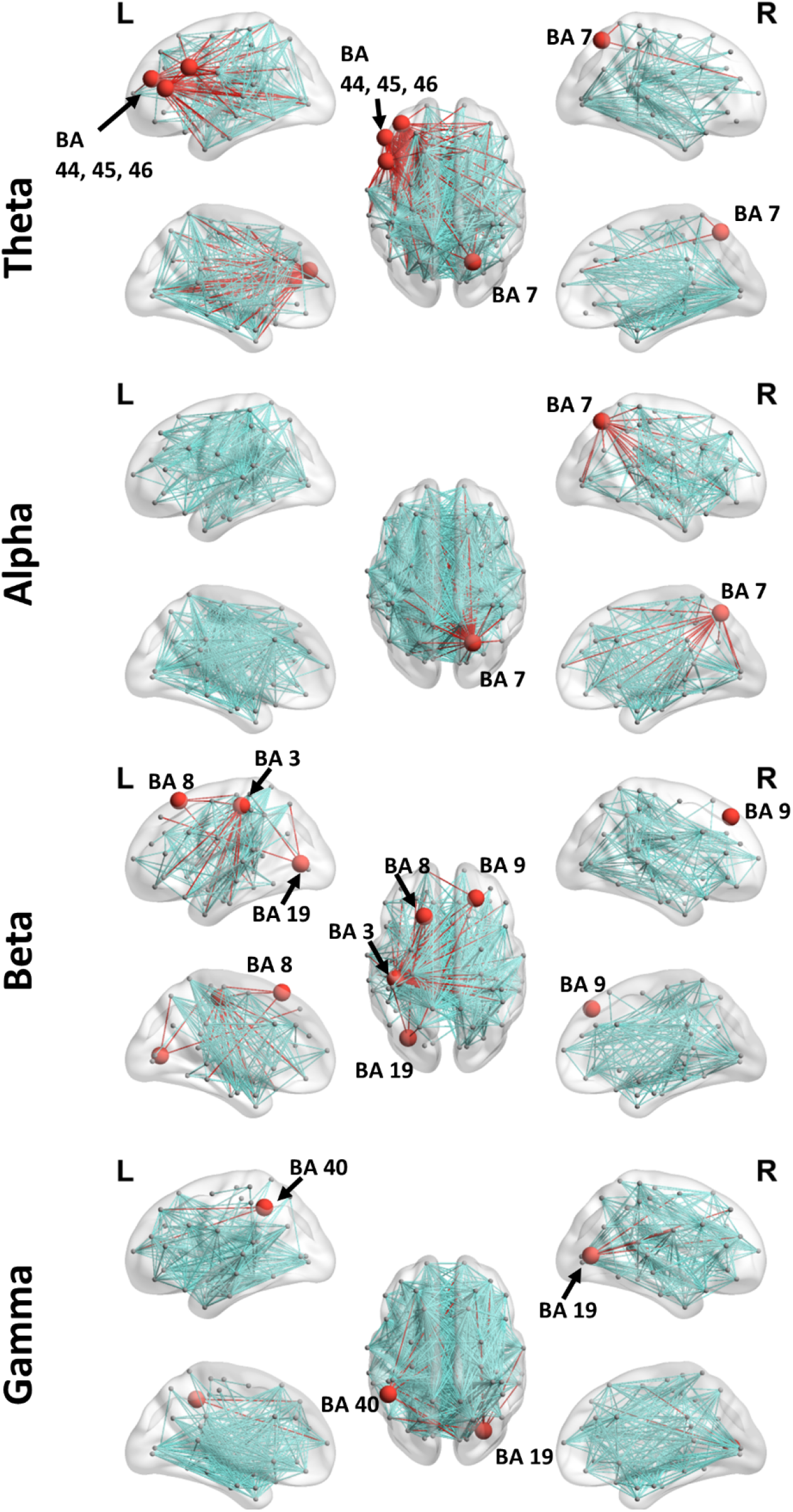
GTA connectivity maps in the *saccade* condition vs. *fixation* condition comparison. Data are projected onto lateral (upper row), upper (middle) and medial (lower row) views of the left (L) and right (R) representations of cortex (generated by BrainNet Viewer toolbox(Xia et al. 2013), MALTAB R2019). Brodmann areas that showed significant differences of centrality between conditions were considered as hubs and are shown by bolded red dots, and their connections are highlighted in red. The 20% of strongest connections related to differences between conditions (*saccade-fixation*) are shown as connectivity lines (edges), in blue-green for non-hub connections.

In the theta band, left BAs 44 (t= 3.86, p=0.001), 45 (t= 4.28, p<0.001), 46 (t= 2.9, p=0.01) showed significant differences of *Eig_C_*. These regions include inferior frontal and dorsolateral prefrontal cortex. These regions were densely connected to temporoparietal regions. Furthermore, in this band, a significant difference in *Eig_C_* was also observed in the right BA 7 (t= 3.11, p=006), including precuneus. This region showed functional connectivity with frontoparietal regions. In the alpha band, significant differences of *Eig_C_* were observed in right BA 7 (t=2.31, p=0.033). This region exhibited connectivity with frontoparietal regions. In the Beta band, significant differences in *Eig_c_* were found in left BAs 3 (t=2.83, p=0.012), 8 (t=3.48, p=0.003), 19 (t=3.74, p=0.002) (a part of visual cortex) and right BA 9 (t=3.14, p=0.006) which is part of the medial and dorsolateral prefrontal cortex. In this band the connectivity pattern was complex, with widely connections from these areas to other brain regions. Finally, in the gamma band, significant differences in *Eig_C_* were observed in left Brodmann 40 (t= 2.55, 0.021) (which includes SMG) and right BA 19 (t=2.80, p=0.003) which is a part of the occipital lobe. In this band left SMG (BA 40) exhibited connectivity to frontal and occipital regions and right visual cortex showed connectivity to temporal and frontal regions.

Figure 9 shows the distribution of each network measure across individuals in different bands for the interaction [(SR-S) – (FR-F)] comparison, using the same conventions as Figure 7. Here, between SR-S and FR-F, the clustering coefficient *(CC), Global Efficiency (Ef), Energy H, and entropy (S)* were significantly different. In the *SR-S*, the clustering coefficient and global efficiency were higher, and energy and entropy were lower in comparison to the *FR-F* (see Figure legend 10 for statistical details).

**Figure 9:**
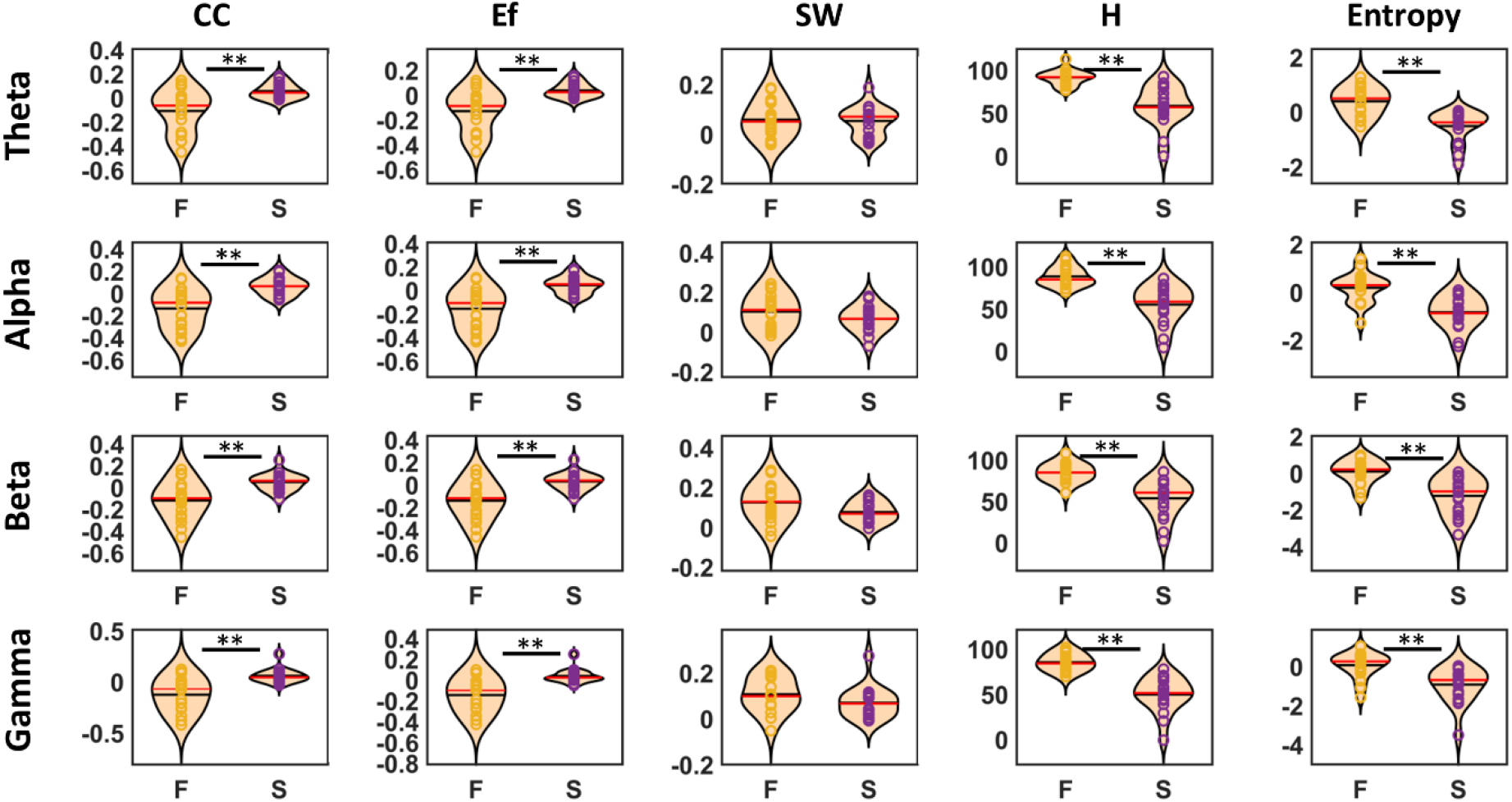
The violin bar plots of network indices in the *fixation* and the *saccade* conditions. Each dot present one individual and the width/shape of violin plots shows distributions of measures for all individuals (Violin plots combine the attributes of standard bar graphs and standard frequency histograms but arranged vertically with frequency shown bidirectionally in the horizontal dimension). The * indicates p<0.05 and the ** indicates p<0.005. F: fixation condition, S: saccade condition, CC: clustering coefficient, Ef: global efficiency, SW: smallworldness, and H: energy. Statistical details: *Clustering* coefficient (theta: t= 3.59, p= 0.002; alpha: t= 4.92, p<0.001; beta: t= 4.18, p<0.001; gamma: t= 4.62, p<0.001), *global efficiency* (theta: t=3.85, p=0.001; alpha: t= 5.23, p<0.001; beta: t= 4.43, p<0.001; gamma: t= 4.79, p<0.001), *energy* (theta: t= 6.20, p<0.001; alpha: t= 5.47, p<0.001; beta: t= 5.10, p<0.001; gamma: t= 8.05, p<0.001), *entropy* (theta: t= 4.33, p= p<0.001; alpha: t= 4.66, p<0.001; beta: t= 4.14, p<0.001; gamma: t= 3.99, p<0.001)

#### 3-2-3 Post hoc analysis: influence of stimulus repetition in the presaccadic interval

A post hoc analysis was performed for separate trials in the *saccade* and *fixation* tasks. After removing the saccade data, no significant differences remained between *Eig_C_* values in the *Fixation* condition and *fixation/repetition* condition. However, several significant differences were observed between the *saccade/repetition* and the *saccade* conditions in the network features. Figure 10 shows these differences using the same analysis and conventions as Figures 6 and 8. In the Theta band, *Eig_C_* was significantly repetition-dependent in the right BA 7 (t=3.25, p=0.004) which includes precuneus, with connections to frontoparietal cortex. In the alpha band, right BA 39 (t=3.09, t=0.007), containing angular gyrus and SMG, also connected to frontoparietal cortex. In the gamma band, several significant hubs showed up, including left BA 40 / SMG (t=2.42, t=0.027), right BA 46 / dorsolateral prefrontal cortex (t=0.013, p=0.013), area 17 / primary visual cortex. etc. Each of these areas showed diffuse functional connectivity in this band. Overall, this comparison engaged areas related to low-level vision, high-level vision, and saccades, with widespread functional connectivity throughout cortex.

**Figure 10:**
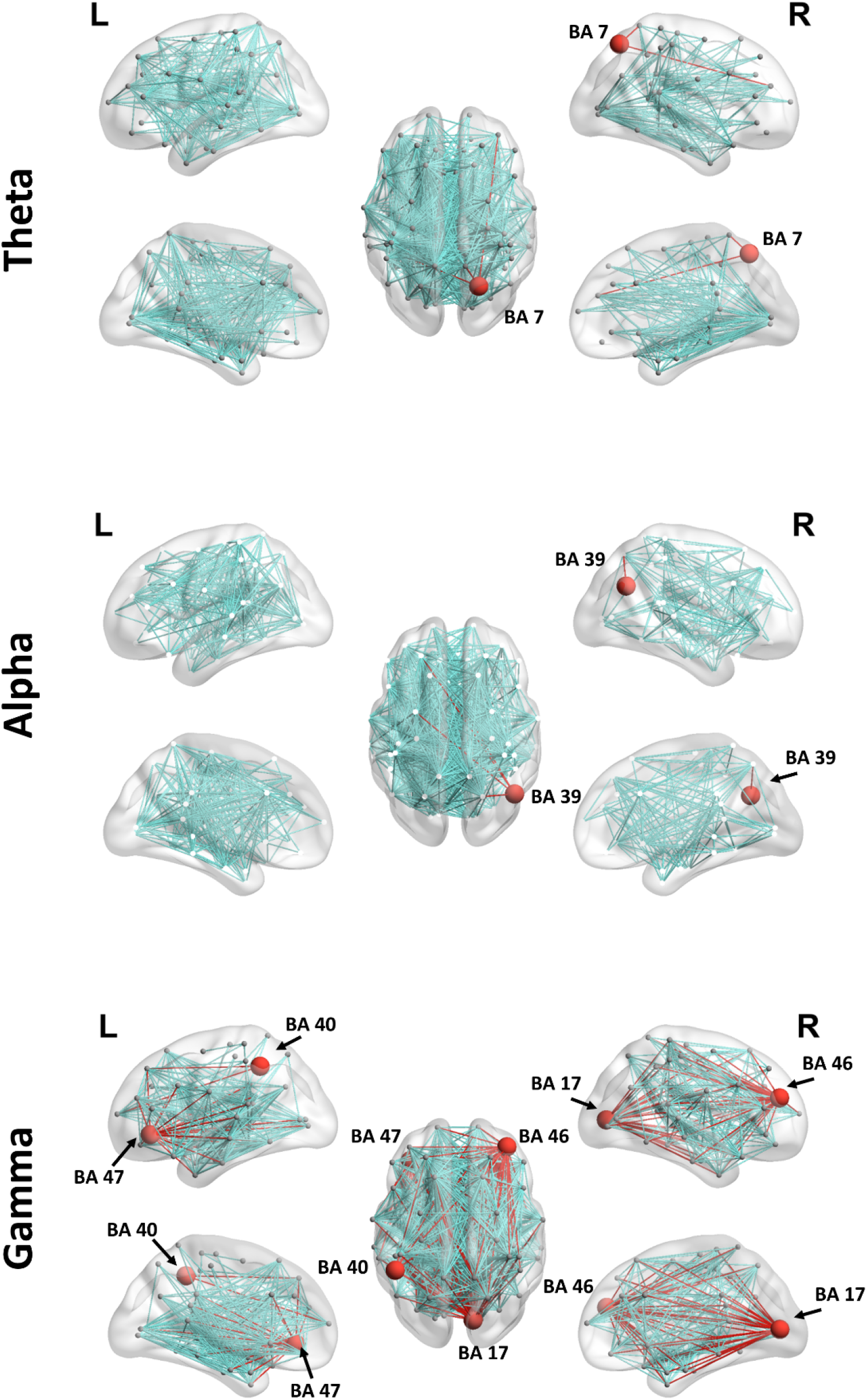
GTA connectivity maps in the *saccade* condition vs. *fixation* condition comparison. Data are projected onto lateral (upper row), upper (middle) and medial (lower row) views of the left (L) and right (R) representations of cortex (generated by BrainNet Viewer toolbox(Xia et al. 2013), MALTAB R2019). Brodmann areas that showed significant differences of centrality between conditions were considered as hubs and are shown by bolded red dots, and their connections are highlighted in red. The 20% of strongest connections related to differences between conditions (*saccade-fixation*) are shown as connectivity lines (edges), in blue-green for non-hub connections.

Figure 11 shows the graph measure distributions the SR-S Comparison, using similar conventions to Figures 7 and 9. In the alpha band, the entropy was significantly lower in the *saccade/repetition* condition in comparison to the *saccade* condition (t=2.34, p=0.032) that means the complexity of functional brain network was decreased in the *saccade/repetition* condition. In the Beta band, the *CC* (t=2.35, p=0.031), and the *Ef* (t=2.34, p=0.032) were significantly higher in the *saccade/repetition* condition while the *S* was significantly higher (t=2.72, p=0.014) in the *saccade* condition. This suggests increased integration and segregation but decreased complexity in the *saccade/rep*etition condition. In the gamma band, the results were similar to the alpha band and *S* was significantly higher in the *saccade* condition (t=2.09, p=0.040) in comparison to the *saccade/repetition* condition. This result also suggests decreased complexity in the *saccade/rep*etition condition. Briefly, these results reveal that repetition of a pre-saccadic stimulus can increase the segregation and integration of functional brain networks but decreases complexity in these networks.

**Figure 11:**
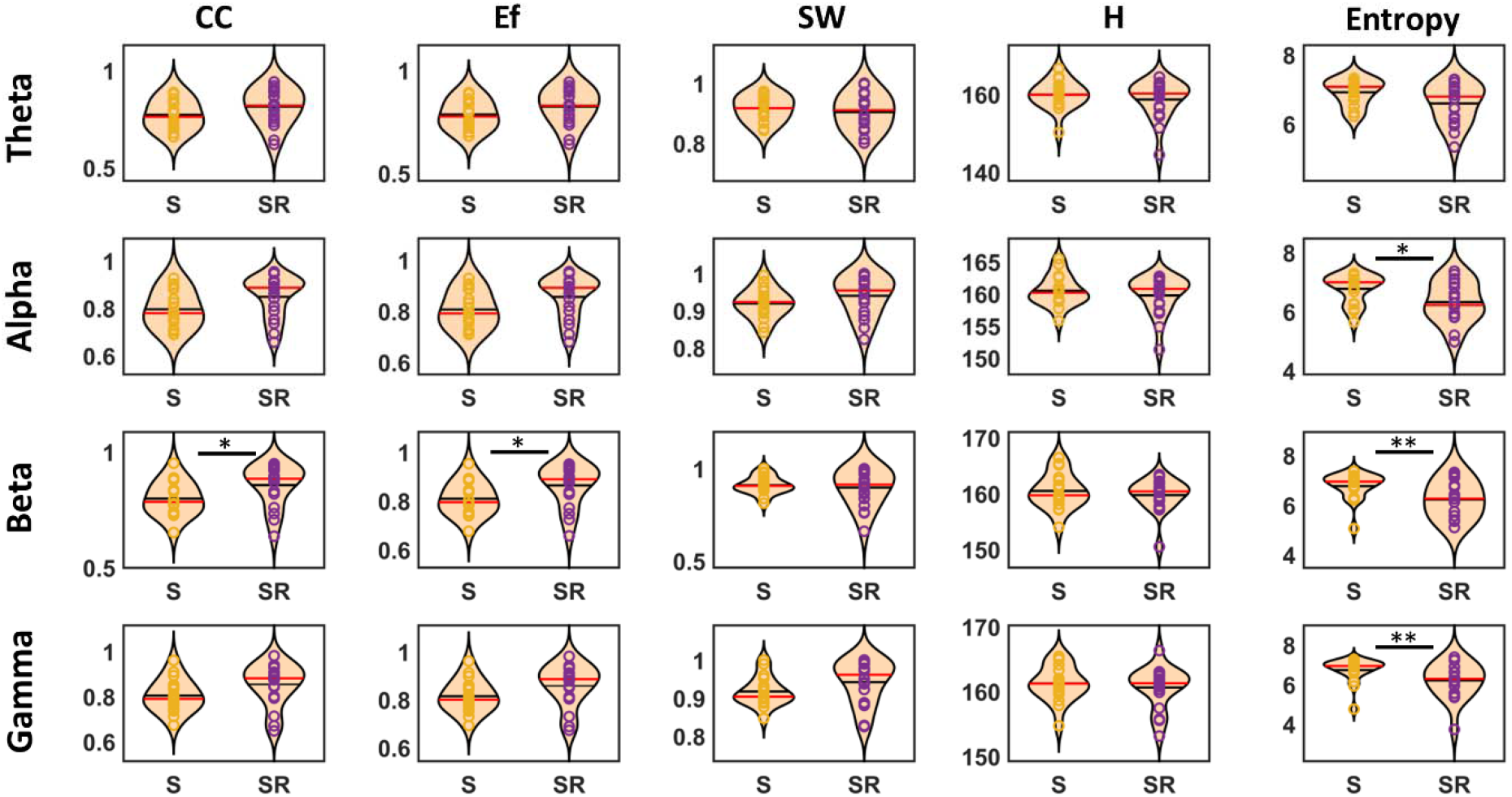
The violin bar plots of network indices in the *fixation* and the *saccade* conditions. Each dot present one individual and the width/shape of violin plots shows distributions of measures for all individuals (Violin plots combine the attributes of standard bar graphs and standard frequency histograms but arranged vertically with frequency shown bidirectionally in the horizontal dimension). The * indicates p<0.05 and the ** indicates p<0.005. F: fixation condition, S: saccade condition, CC: clustering coefficient, Ef: global efficiency, SW: small-worldness, and H: energy.

## 4 Discussion

EEG / MEG have been employed previously to identify fast neural responses in the presaccadic interval (Forgacs et al. 2008; Jerbi et al. 2009; Kern, Schulze-Bonhage, and Ball 2021; Kovalenko and Busch 2016; McLelland et al. 2016; Medendorp et al. 2007), but this study introduced several innovations: first, the addition of pre-saccadic stimulus repetition to probe saccade-vision interactions, second, the use of EEG source localization, and third the use of GTA to quantify functional bran network topology/dynamics. Overall, our results show that during the presaccadic interval, saccades caused modulations in dorsomedial parietofrontal cortex, with decreased current density in all four frequency bands (theta, alpha, beta, gamma) and prominent connectivity in BA 8 (frontal eye fields) and 39 (inferior parietal cortex) in the alpha and beta bands. Presaccadic stimulus repetition interacted with the saccade-fixation at both local and network levels, with increased bilateral precuneus current density and highly increased connectivity for a number of band-specific hubs. Post-hoc analysis revealed that this was primarily due to presaccadic interactions with motor signals in the saccade condition, where repetition decreased frontoparietal current density in the alpha and beta bands and increased network connectivity in various hubs. Importantly, saccades and stimulus repetition both produced enhanced network segregation and integration, although they had opposite effects on entropy. In the following sections we discuss and interpret these results in more detail.

### 4-1 Functional anatomy of presaccadic signals in frontoparietal cortex

Previous neurophysiological and neuroimaging studies have implicated superior and medial frontal cortex (specifically FEF/SEF respectively) and parietal cortex (especially monkey intraparietal cortex) in saccade planning and remapping ((Anderson et al., 1994; Barash et al., 1991; Schall & Hanes, 1993; Schall et al., 1995; Umeno & Goldberg, 1997)). Our source localization analysis showed reduced current density (in all frequency bands) in dorsomedial frontal cortex and the superior parietal lobule. Note that reduced current density has been associated with desynchronization, and thus greater complexity of neural activity (Blohm et al. 2019). These results are consistent with previous studies showing alpha and beta desynchronization in superior/medial frontal regions and superior parietal cortex during saccades and other actions (Brignani et al. 2007; Leocani et al.; 1997; Stancák & Pfurtscheller, 1996; Tzagarakis et al. 2010).

Prefrontal regions of presaccadic current density modulation included 1) the classic frontal eye fields (Anderson et al. 1994; Schall and Hanes 1993), 2) more medial areas that could contain dorsolateral prefrontal cortex (DLPFC), a saccade-modulated area involved in working memory and executive functions (Balan & Ferrera, 2003; Takeda & Funahashi, 2002). We also observed altered presaccadic current density in the medial posterior parietal cortex (precuneus). Precuneus is not considered part of the classical oculomotor system, but is activated in tasks that involve perisaccadic processing of multiple, complex visual stimuli (Chen et al. 2018; Frings et al. 2006). Precuneus activation may have been associated with additional demands in our particular task, such as the increased difficulty of judging stimulus duration in the presaccadic interval (Gitelman et al., 1999; Huddleston et al., 2021; Mort et al., 2003).

Likewise, our network analysis identified major presaccadic ‘hubs’ (i.e., sites that show a high degree of functional connectivity with other well-connected sites) in bilateral BA 8 (in the beta band) and left BA 39 (in the alpha band). BA 8 encompasses the FEF, and BA 39 includes the regions of posterior parietal and especially angular gyrus. Angular gyrus has previously been implicated in saccade memory / planning and may be part of the human expansion of the monkey parietal eye fields (Vesia & Crawford, 2012; Vesia et al., 2010). These results generally overlapped with our source localization results, but whereas the latter included more medial peaks (associated with ‘cognitive’ areas), GTA tended to highlight more lateral sites associated with saccade production (Vesia and Crawford 2012). This could arise because current densities are driven by local processing whereas connectivity analysis is more influenced by longer-range transmission of motor signals. It should also be noted that these results were frequency band dependent. For example, medial parieto-frontal current densities showed strongly in theta and gamma bands associated with memory (Kawasaki & Yamaguchi, 2013; Pesaran et al., 2002), whereas presaccadic hubs showed best in alpha and beta bands associated with sensorimotor activity (Buchholzet al., 2014).

Overall, our source localization and GTA ‘hub’ results showed good agreement both with each other and previous neurophysiological, neuroimaging, stimulation, and neuropsychological data, with the exceptions noted above. Taken together, our source and GTA results confirm that human frontal and parietal eye fields show both greater activity and connectivity just before a saccade, consistent with their known role in saccade generation and high-speed neural signal transmissions in presaccadic interval.

### 4-2 Functional network features in the presaccadic interval

The presaccadic network hubs described above showed extensive connectivity throughout the brain, with major ipsilateral connections in the beta band from Brodmann area 8 (FEF) to superior posterior parietal cortex and other frontal regions (Figure 6). Brodmann area 39 (inferior parietal cortex) showed extensive bilateral connectivity in the alpha band with various frontal lobe regions implicated in saccade processing (Anderson et al., 1994; Funahashi et al., 1991) as well as contralateral parietal cortex. Although functional connectivity does not necessarily imply direct anatomic connection, this is consistent with the known anatomic connections between parietal and frontal saccade areas, and likely involves both feedforward and recurrent processing of saccade planning signals (Alkan et al., 2011; Andersen et al., 1990; Anderson et al., 1994; Corbetta et al., 1998).

From the functional brain network prospective, the afore mentioned regions fall within the overlapping cortical space of two well-established networks: the frontoparietal control and dorsal attention networks (Corbetta and Shulman 2002; Spreng et al. 2010). Frontoparietal control network (encompasses prefrontal cortex including FEF, lateral intraparietal sulcus, superior parietal regions, and dorsal precuneus regions) is involved in cognitive control and working memory (Niendam et al. 2012). Previous investigations have reported that saccades alter activity the frontoparietal control network, including the regions reported here (Scolari, Seidl-Rathkopf, and Kastner 2015). The same regions (but with different patterns of activation and connectivity) are involved in the dorsal attention network (Szczepanski et al. 2013). This network is involved in orienting attention toward a specific target, an important aspect of saccade production (Basso and Wurtz 1997).

Besides providing measures of specific connectivity, GTA analysis provides global quantitative indices of functional network topology, such as clustering coefficient (*CC*), and efficiency (*Ef*), all of which increased here in the period before saccades. The *CC* and the *Ef* in the functional brain networks are associated with network segregation and network integration, respectively (Bullmore & Sporns, 2009; Ghaderi et al., 2017, 2018b). Higher values of the *CC* exhibit higher level of information processing in the local circuits whereas higher values of the *Ef* indicate faster information propagation in the large scale brain network (Rubinov and Sporns 2010). Networks with high values of the *CC* and the *Ef* are efficient networks for information transformation in the local and global levels (Watts & Strogatz, 1998; Rubinov & Sporns, 2010). Thus, network segregation and integration increased in the presaccadic interval, suggesting enhanced global network integrate information across separate cortical modules.

Furthermore, our results showed that the energy (*H*) and Shannon entropy (*S*) in the functional brain network are also increased during the presaccadic interval. The *H* value is related to stability of synchronization in the functional brain network (Daianu et al., 2015; Ghaderi et al., 2020, 2021) and *S* is associated with complexity of network (Ghaderi et al., 2019, 2020, 2021). This result suggests that synchronization between functional network nodes (all brain regions) is more stable during the presaccadic interval, which can support fast propagation between cortical nodes. Further, this was accompanied by increased *S* values, suggesting increased complexity of information processing in the brain. This is not a trivial observation: in some functional brain networks *H* and *S* show inverse correlations (Ghaderi et al. 2020). Since the value of *S* in a complex network is related to diversity of functional weights, the increased *S* in the presaccadic interval shows the brain uses more diverse neural pathways for information processing in the presaccadic interval.

Since the segregation, integration, synchronization, and complexity of brain network were simultaneously and specifically enhanced during the presaccadic interval (compared to fixation), one can justify the existence of a *presaccadic* network. This highly efficient and synchronized network could provide a neural basis for the rapid integration of pre- and post-saccadic visual information during the saccade.

Conversely, recent evidence suggests that time, space and movement perception are closely related to states of transient neural network, then the *presaccadic* network can be an appropriate candidate to explain various perisaccadic distortions in time, space and movement (Burr et al. 2011; Eagleman 2005). For example, the features of the *presaccadic* network can be associated with saccadic *remapping* that needs very fast information processing on the local level and rapid integration between different brain areas on the global level (Morrone et al. 2005). This suggests that the presaccadic network should interact with presaccadic visual signals, as we consider next.

### 4-3 Stimulus repetition alters the presaccadic cortical network

We employed presaccadic stimulus repetition as a means to probe signal interactions between visual processing and saccades at both local and network levels. Previous psychophysical studies have indicated that stimulus repetition influences perception of a subsequently presented novel stimulus (Grill-Spector et al. 2006; Matthews and Gheorghiu 2016), but here it yielded only modest main EEG effects in our combined saccade-fixation dataset. However, stimulus repetition yielded significant interactions with fixation vs. saccades. This interaction was accompanied by an alteration in both current density (local activation) and centrality (network connectivity) in precuneus, again consistent with a role for this structure precuneus in complex task-related visual processing in the presaccadic interval (Dyckman et al., 2007; Mort et al., 2003; Simon et al., 2002). Further, this interaction affected the centrality of dorsolateral prefrontal cortex. Lateral prefrontal cortex is involved in executive control for saccade tasks (Koval et al., 2011; Ploner et al., 2005) and increased attention to a novel visual stimulus (Heeman et al., 2014). Lastly, SMG centrality was altered by saccade-repetition interactions, consistent with recent fMRI studies that showed a specific sensitivity of SMG to stimulus orientation changes (similar to those here) in the presence of saccades (Dunkley et al., 2016; Baltaretu et al. 2020).

When we performed a post-hoc analysis on fixation and saccades separately, stimulus repetition had no significant effect in our EEG blocks of fixation data. This is consistent with previous EEG/MEG studies showing no significant effects of repetition over short time scales (<100ms) after novel stimulus onset (Cycowicz & Friedman, 2007; Friedman, Cycowicz, & Gaeta, 2001). However, when a repeated series of stimulus was presented before the saccade (saccade/repetition condition), the regional current density and network features in the perisaccadic interval were significantly influenced by stimulus repetition. In lower frequencies (i.e., theta and alpha bands), presaccadic stimulus repetition decreased current density in frontal and parietal cortex, whereas in the beta band current density increased in the central areas, and in the gamma band, current density is increased in posterior regions while decreasing in the frontal cortex.

Furthermore, right superior parietal lobe / precuneus (theta band), right inferior parietal lobe (alpha band), and V1 / dorsolateral prefrontal cortex (gamma band) exhibited higher *Eig_C_* when a pre-saccadic stimulus was repeatedly presented before saccade, signifying more involvement in a functional network (Fig. 10). On the other hand, SMG (gamma band) exhibited decreased *Eig_C_* during presaccadic repetition perhaps consistent with repetition suppression seen in SMG during fMRI studies signals (Baltaretu et al., 2020; Dunkley et al., 2016). This area may be a human extension of the lateral intraparietal cortex in the monkey (Subramanian and Colby 2014) and plays a role in trans-saccadic integration of visual signals (Baltaretu et al. 2020; Dunkley et al. 2016). These various hubs, in turn, showed widespread connectivity throughout occipital, temporal, parietal, and frontal cortex, indicating extensive saccade-dependent sharing of visual signals throughout the brain.

Again, from the functional network perspective, the above mentioned areas are consistent with the frontoparietal control and dorsal attention networks. As noted above (section 4-2), these networks are involved in saccades and visual attention, and they have also been implicated in the processing of signal repetition (Kim 2017). Therefore, it makes sense that we find them involved in saccade-repetition interactions. These frontoparietal interactions might be related to increased cognitive control demands, signal noise, and the taxing of visual working memory during saccade planning (Finc et al. 2020; Prime et al. 2007; Sajad et al. 2016). This result (overlapping of involved networks in presaccadic and repetition signals) might be responsible for the subjective perceptual phenomena observed in our behavioral study (Ghaderi, Niemeier, and Crawford, 2021), a topic that requires further investigation.

Overall, our GTA results showed that the pre-saccadic repetition reduced the entropy of presaccadic network in the theta, beta, and gamma bands. However, clustering coefficient and global efficiency was increased after pre-saccadic repetition. This decreased complexity and increased integration and segregation with same level of synchronizability (for the same energy), suggests the level of information processing was reduced in the presaccadic interval when a pre-saccadic stimulus was presented repeatedly before the saccade. However, this processing was occurring in a more topologically efficient network (with higher integration and segregation). Based on this, one can say that presaccadic stimulus repetition facilitates information processing in the presaccadic network. More generally, this suggests that in the presaccadic interval, visual processing networks are extensively influenced throughout the brain. These altered network properties may also be associated with perisaccadic changes in space and/or time perception (Burr et al. 2011; Eagleman 2005; Morrone et al. 2005).

## 5 Conclusion

The results of this study reveal new aspects of neural mechanisms in the presaccadic interval. First, the use of high-density EEG in combination with source localization largely confirms the regional cortical specificity for saccades and presaccadic stimulus repetition observed in fMRI studies, but over more specific time scales (and broader accessibility than MEG). More importantly, the application of Graph Theory Analysis provided objective identification of hubs and functional networks for these processes within a relatively large-scale presaccadic network. Finally, topological analysis of this presaccadic network showed increased levels of integration, segregation, synchronizability and complexity, indicating an accelerated rate and scale of cortical information processing just before saccades. Overall, this study identifies a *presaccadic network* with major hubs in parieto-frontal cortex, but with connectivity and interactions / influence on visual processing throughout the cerebral cortex. It is likely that this approach will prove useful in explaining the various perceptual distortions that occur in the presaccadic interval.

## Acknowledgements

We thank Dr. X Yan for technical support. This research was funded by the National Science and Engineering Research Council of Canada (NSERC). J.D. Crawford was supported by a Canada Research Chair. A. Ghaderi held a Vision: Science to Application (VISTA) Fellowship, supported by the Canada First Research Excellence Fund. M. Niemeier was supported by an NSERC Discovery grant.

